# Src/Fas2-dependent Ephrin phosphorylation initiates Eph/Ephrin reverse signaling through Rac1 to shape columnar units in the fly brain

**DOI:** 10.1101/2023.09.24.559153

**Authors:** Miaoxing Wang, Xujun Han, Yunfei Lee, Rie Takayama, Makoto Sato

## Abstract

Columns are the morphological and functional units containing multiple neurons in the brain. The molecular mechanisms of column formation are largely unknown. Ephrin/Eph signaling mediates a variety of developmental processes. Ephrin acts as a ligand for Eph to regulate forward signaling, while Eph acts as a ligand for Ephrin to regulate reverse signaling. However, whether and how the uni- or bi-directional Ephrin/Eph signaling is involved in column formation remains elusive. In this study, we show that Ephrin and Eph regulate the morphology and location of columnar neurons through bi-directional repulsive signaling. Furthermore, Eph ligand triggers cytoplasmic tyrosine phosphorylation of Ephrin under the control of Src kinases and Fasciclin II (Fas2), forming Ephrin/Src/Fas2 complex to promote reverse signaling through a downstream regulator, Rac1. This study presents for the first time a unified picture of the molecular interactions in the defined context of column formation using the fly brain as a model.

## Introduction

In various brain tissues, columns are the structural and functional units containing multiple neurons. Within a column, the morphology and projection of diverse neurons are highly organized to form precise neural circuits. However, the developmental mechanism of columnar unit formation remains largely unknown.

The brain of *Drosophila melanogaster* is an excellent model for exploring the genetic and molecular mechanisms of column formation, as it shares similar columnar structures with the mammalian brain with fewer neurons included (*1–3*). The fly visual system includes the retina and the optic lobe, which consists of the lamina, medulla, and lobula complex. The medulla, the largest neuropil of the optic lobe, contains approximately 100 morphologically distinct types of neurons forming 800 columns (*4*). Visual information received by photoreceptors in the retina is relayed through the lamina to the distal medulla. R7 and R8 photoreceptors and L1–L5 lamina neurons directly innervate the medulla neuropil, and it is known that the terminals of R7, R8, and L1–L5 neurons compose a single medulla column together with the other columnar medulla neurons (*5*). The medulla intrinsic neuron Mi1 is a uni-columnar neuron, projecting to the peripheral edge region of the columns during larval development. Our previous study demonstrated that R7, R8, and Mi1 are the core columnar neurons that are concentrically arranged in the larval medulla according to N-cadherin (Ncad)-dependent differential adhesion (*3*). The terminal of R7 occupies the dot-like central region of the column. The R8 terminal enwraps the R7 terminal forming a donut-like region, and the Mi1 terminal occupies a grid-like region outside the R8 terminal (Figs. 1A-D). The precise organization of these columnar neurons is regulated by multiple molecules (*3, 6–8*).

**Fig. 1.**
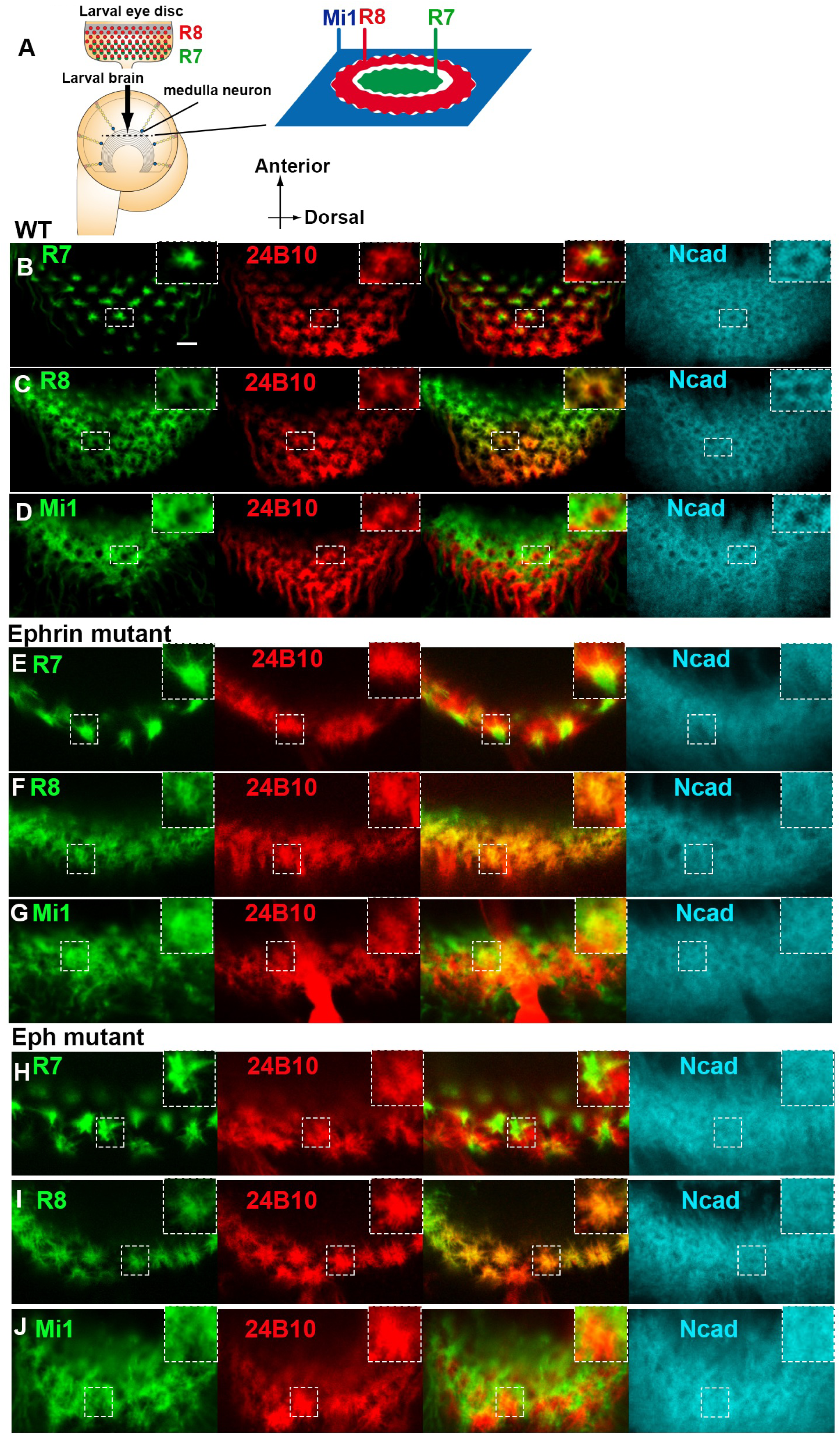
Ephrin and Eph regulate the morphology of neurites of columnar neurons. **A** Schematics of the developing larval visual systems. In the larval eye disc, R8 and R7 are sequentially differentiated behind the morphogenetic furrow. Columns are identifiable along the planes indicated by dotted lines in the optic lobe. (**A-D**) R7 terminal (**B**; green) occupies the dot-like central region of the column shown by Ncad (blue), R8 terminal (**C**; green) enwraps R7 terminal forming a donut-like region, and Mi1 terminal (**D**; green) occupies a grid-like region outside the R8 terminal. 24B10 (red) visualizes R8 terminals. (**E-J**) In *Ephrin*-null mutant (*Ephrin^I95^*) (**E-G**) and *Eph*-null mutant (*Eph^X652^*) (**H-J**), the morphology of the three core columnar neurons is disorganized: R7 shows an expanded axon terminal; R8 axon terminal penetrates the central hole region disrupting the donut-like pattern; Mi1 neurite also penetrates the central hole region disrupting the grid-like pattern. Scale bar, 5 μm.

Ephrin and Eph have been shown to be involved in not only diverse developmental processes but also a variety of diseases (*9–12*). They can both act as ligand and receptor for forward and reverse signaling through cell-to-cell communication (*13–16*). In vertebrates, they are divided into two main subfamilies of ligand/receptor couples, Ephrin-A/Eph-A and Ephrin-B/Eph-B, based on their specific structures (*17, 18*). Ephrin-B reverse signaling has been shown to have multiple functions in various developmental processes that are dependent on or independent from its cytoplasmic tyrosine phosphorylation. Three out of five tyrosines conserved in the Ephrin-B subfamily have been characterized as the major phosphorylation sites *in vitro* (*19*). It has been reported that Src family kinases are positive regulators of Ephrin-B phosphorylation (*20*). However, these studies are essentially based on cultured cells and/or *in vitro* systems. The molecular mechanism and biological significance of Ephrin phosphorylation *in vivo* still remain elusive.

Ephrin-A and Ephrin-B have been shown to be involved in column formation by integrating the distribution of cortical pyramidal neurons at a macroscopic level (*21, 22*). In the fly visual system, the Ephrin homologue related to Ephrin-B is required for topographic map formation and branching restriction of lamina neurons (*23–25*). However, whether and how the uni- or bi-directional Ephrin/Eph signaling is involved in column formation is elusive.

In this study, we demonstrate that Ephrin and Eph regulate the morphology and location of core columnar neurons during column formation through bi-directional repulsive signaling *in vivo* using the fly brain as a model. We generated Ephrin antibodies that specifically detect phosphorylated and un-phosphorylated forms of its intracellular domain. Using these antibodies, we found that they show different distribution patterns complementary with each other in the core columnar neurons. Phosphorylated Ephrin is specifically localized in R7 showing a dot-like pattern while un-phosphorylated Ephrin and Eph are localized more widely in R8 and Mil showing a grid-like expression pattern. They regulate the morphology and segregation of the core neuron terminals by the bi-directional repulsive activity of Ephrin/Eph-signaling through interaction between the columnar neurons. Ephrin expressed in R7 is phosphorylated on tyrosine residues in the cytoplasmic domain and is regulated by Src kinases and the *Drosophila* orthologue of neural cell adhesion molecule (NCAM), Fasciclin2 (Fas2), under the control of Eph expressed in R8. Furthermore, we also found that Rac1 acts as a downstream effector of Eph/Ephrin reverse signaling. Thus, this study presented for the first time a unified picture of the complex molecular interactions in the defined context of column formation.

## Results

### Ephrin/Eph signaling is required for columnar neuron organization

To investigate how *Ephrin* and *Eph* function in column formation, we first observed the morphology and organization of the core columnar neurons in *Ephrin* and *Eph*-null mutants. Since column formation is initiated by three core neurons, R7, R8, and Mi1, establishing distinct concentric domains within a column (Fig. 1A), we focus on the larval medulla as a model system to examine the roles of *Ephrin* and *Eph* in column formation. In the wild type, the terminal of R7 occupies the dot-like central region of the column, the R8 terminal enwraps the R7 terminal forming a donut-like region, and the Mi1 terminal occupies a grid-like region outside the R8 terminal (Figs. 1A-D). In the *Ephrin*-null mutant (*Ephrin^I95^*), the morphology of all these three core columnar neurons was disorganized: R7 shows an expanded axon terminal; R8 axon terminal penetrates the central hole region disrupting the donut-like pattern; Mi1 neurite also penetrates the central hole region disrupting the grid-like pattern (Figs. 1E-G). A similar phenotype was observed in the *Eph*-null mutant (*Eph^X652^*; Figs. 1H-J), suggesting that both *Ephrin* and *Eph* are involved in organization of the columnar neurons. The number of R7, R8 and Mi1 terminals was decreased in this and following experiments most likely because their projections to the medulla neuropil are also regulated by Ephrin and Eph. However, we solely focus on the interaction between their terminals in the medulla neuropil in this study.

### Phosphorylated and un-phosphorylated Ephrin show distinct distribution patterns

Next, we investigated the expression patterns of Ephrin and Eph in the developing medulla. The expression of Eph was visualized using the *Myc* knock-in allele of *Eph* (*26*). Anti-Myc antibody staining revealed a grid-like expression of Eph-Myc at the late third instar larval stage (Fig. 2A) and a donut-like pattern at 24h and 48h after puparium formation (APF) overlapping with Ncad staining (Figs. S1A, S1B). To visualize Ephrin distribution, we generated antibodies that recognize the intracellular domain of *Drosophila* Ephrin (Fig. 2F). Interestingly, the two Ephrin antibodies named Ephrin #1 and Ephrin #2 show different patterns that are complementary with each other. Ephrin #1 shows a dot-like pattern similar to the pattern of R7 terminals throughout development (Figs. 2B, S1C, S1D). In contrast, Ephrin #2 shows a grid-like pattern similar to the pattern of Eph-Myc expression (Figs. 2A, 2C, S1E, S1F). In *Ephrin* mutant brains, the signals of Ephrin #1 and Ephrin #2 were lost, suggesting that both antibodies recognize Ephrin protein (Figs. 2D, 2E).

**Fig. 2.**
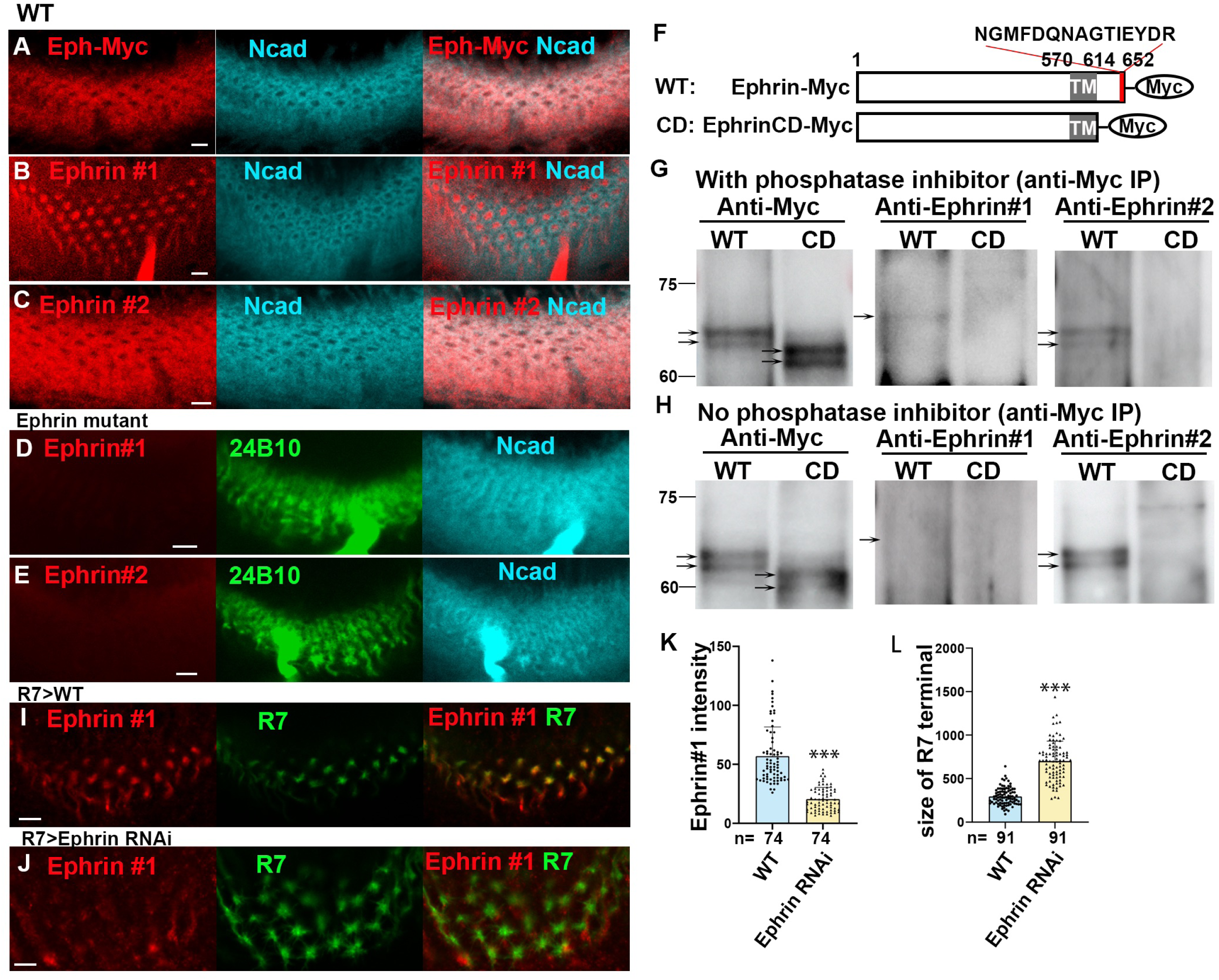
Distributions of Eph and Ephrin in columnar neurons. (**A**) Anti-Myc antibody staining revealed a grid-like expression of Eph-Myc (red) overlapping with Ncad staining (blue). (**B, C**) Ephrin #1 and Ephrin #2 antibodies (red) show different patterns that are complementary to each other. Ephrin #1 shows a dot-like pattern, and Ephrin #2 shows a grid-like pattern. Ncad (blue). (**D, E**) Both the signals of Ephrin #1 and Ephrin #2 were lost in *Ephrin* mutant. (**F**) The structure of Ephrin-Myc (WT) and EphrinCD-Myc (CD), a cytoplasmic deletion mutant. (**G, H**) Validation of Ephrin antibodies by immunoblotting followed by anti-Myc immunoprecipitation. In the presence of phosphatase inhibitor, Ephrin #1 detects one band in WT, while Ephrin #2 detects two bands that are also detected with Myc-antibody in WT (**G**). In the absence of the phosphatase inhibitor, no band is detected by Ephrin #1, while signals of Ephrin #2 are enhanced (**H**). (**I, J**) Knocking down *Ephrin* in R7 decreased Ephrin #1 signals. (**K, L**) Quantification of the intensity of Ephrin #1 signals and the size of R7 terminals (in **I, J**). Results were statistically analyzed using Welch’s t-test (*** indicates p< 0.001). Scale bars, 5 μm.

We further tested the specificity of these two antibodies through immunoprecipitation of Myc-tagged Ephrin proteins overexpressed in photoreceptor neurons under the control of *GMR-Gal4*. Although the Ephrin antibodies recognized the full-length Myc-tagged Ephrin (Ephrin-Myc), they did not recognize the truncated protein that lacks the cytoplasmic domain (EphrinCD-Myc; Figs. 2F-H), suggesting that they both recognize the cytoplasmic domain of Ephrin. Interestingly, Ephrin-Myc detected by Ephrin #1 was larger than that detected by Ephrin #2 in the presence of phosphatase inhibitor (Fig. 2G). However, the former was lost while the latter was increased in the absence of phosphatase inhibitor, suggesting that Ephrin #1 recognizes the phosphorylated Ephrin while Ephrin #2 recognizes the un-phosphorylated Ephrin (Fig. 2H). The faint band detected by Ephrin #1 is consistent with the fact that the Ephrin #1 signal is found only in a limited area in the brain compared with that of Ephrin #2 (Figs. 2B, 2C, 2G).

Here, we assume that both phosphorylated and un-phosphorylated Ephrin exist in the presence of phosphatase inhibitor resulting in the detection of Ephrin-Myc by Ephrin #1 and #2 (Fig. 2G). In the absence of phosphatase inhibitor, un-phosphorylated Ephrin-Myc recognized by Ephrin #2 may become dominant due to the removal of phosphates from Ephrin (Fig. 2H).

### R7 is the source of phosphorylated Ephrin#1

Ephrin #1 shows a dot-like pattern overlapping with R7 cell bodies in the eye and R7 axons in the lamina (Fig. S2) as well as with R7 terminals in the medulla (Fig. 2I). To investigate the source of phosphorylated Ephrin, we knocked down *Ephrin* specifically in R7 and found that Ephrin #1 signal was downregulated while R7 terminals were enlarged (Figs. 2J-L), suggesting that R7 is the source of phosphorylated Ephrin in the medulla.

Meanwhile, Ephrin #2 shows a grid-like pattern overlapping with R8 and Mi1. Ephrin #2 signal was downregulated when *Ephrin* was knocked-down in R8 and Mi1 (Fig. S3), suggesting that R8 and Mi1 are the source of un-phosphorylated Ephrin. Taken together, Ephrin #1 detects the phosphorylated Ephrin in R7, and Ephrin#2 detects the un-phosphorylated Ephrin in R8 and Mi1 in the medulla.

### Ephrin in R7 is autonomously and non-autonomously required for the organization of the columnar neurons

To investigate the function of *Ephrin* in organization of the column, we specifically knocked down *Ephrin* in R7, R8, and Mil neurons. When *Ephrin* was knocked-down in R7, the morphology of R7 and R8 was affected (Figs. 3A, 3B). The terminals of R7 were enlarged (Figs. 3B, 3C). The terminals of R8 were also enlarged, penetrating the central hole region and the neighboring columns (Fig. 3B). The overlap between R8 and R7 terminals increased (Fig. 3D) and the column structure was severely disorganized as visualized by Ncad (Figs. 3A, 3B). These results suggest that phosphorylated Ephrin in R7 regulates the morphology and segregation of R7 and R8 terminals.

**Fig. 3.**
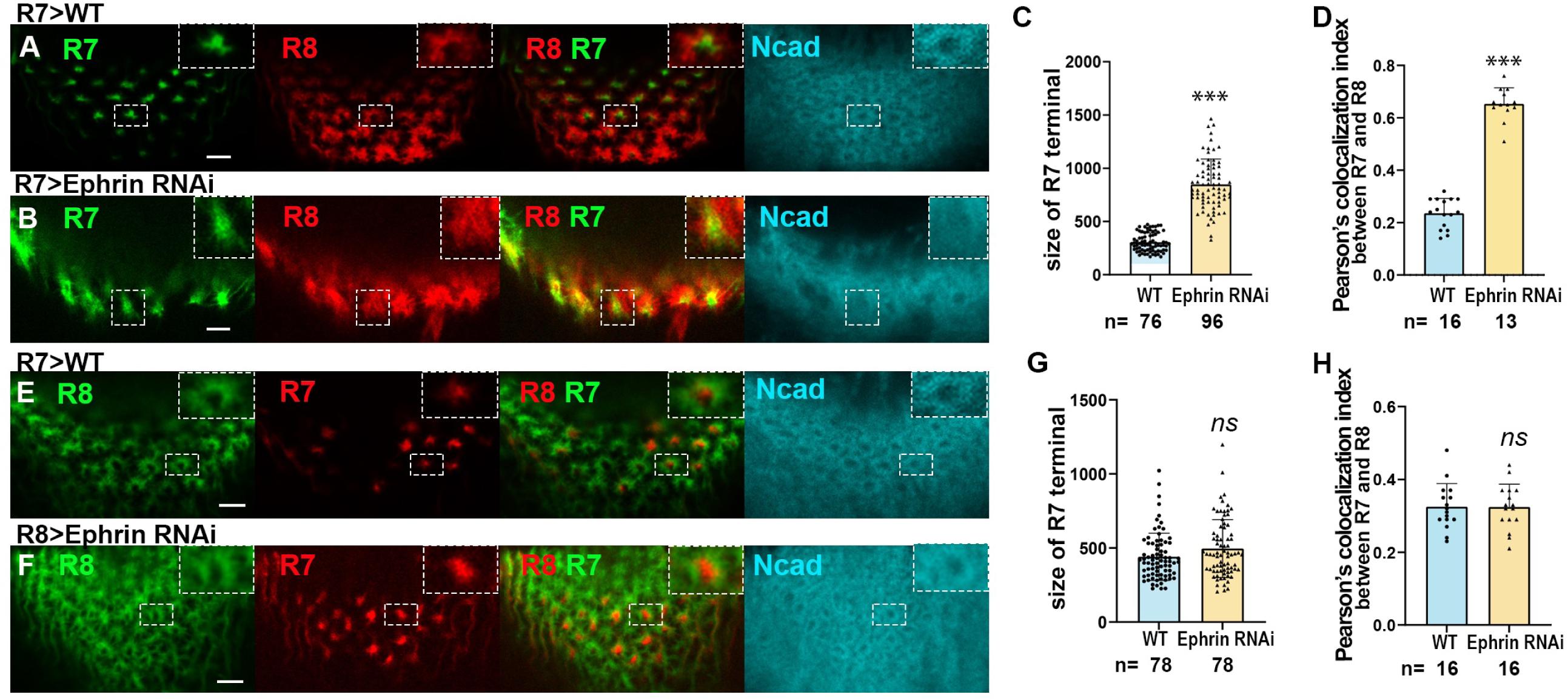
Ephrin expressed in R7 regulates the organization of columnar neurons. (**A, B**) *Ephrin* knock-down in R7 disturbs the separation of R7 (R7-GFP, green) and R8 (R8-RFP, red) terminals and column formation (Ncad, blue). **(C)** Quantification of the size of R7 terminals (in **A, B**). (**D**) Colocalization of the terminals of R7 and R8 (in **A, B**). (**E, F**) *Ephrin* knock-down in R8 did not affect the separation of R7 and R8 terminals and column formation (Ncad, blue). (**G**) Quantification of the size of R7 terminals (in **E, F**). (**H**) Colocalization of the terminals of R7 and R8 (in **E, F**). The dotted boxes are enlarged in the right upper panels (in **A, B, C, F**). Results were statistically analyzed using Welch’s t-test. *** indicates p< 0.001, n.s. not significant. Scale bars, 5 μm.

In contrast, when *Ephrin* was knocked-down in R8, the morphology of R7 and R8 were not significantly affected (Figs. 3E-H). Since the column structure was disrupted compared to the control as visualized by Ncad, other columnar neurons besides R7 and R8 might be affected. Indeed, the morphology of Mi1 was disorganized in the same condition (Figs. S4A, S4B), indicating that un-phosphorylated Ephrin in R8 regulates the morphology of Mi1 terminals.

Since phosphorylated and un-phosphorylated Ephrin are abundant in R7 and R8, respectively, and Eph is strongly localized to R8 and Mi1 neurites, the above results are consistent with the idea that phosphorylated Ephrin in R7 regulates column organization through Eph in R8 and un-phosphorylated Ephrin in R8 regulates column organization through Eph in Mi1.

### Eph in R8 is autonomously and non-autonomously required for the organization of the columnar neurons

Considering the expression pattern of Eph-Myc, Eph is most likely expressed in R8 and/or Mi1 (Fig. 2A). Indeed, *Eph* knockdown in R8 induced morphological defects in R7- and R8-terminals and in column structure as visualized by Ncad (Figs. 4A-D). In contrast, *Eph* knockdown in R7 induced no significant defect (Figs. 4E-H). In addition, *Eph* knockdown in Mi1 induced strong morphological defects in Mi1 while the terminals of R8 were not significantly disrupted (Figs. S4C, S4D), suggesting that Eph is cell autonomously and non-autonomously required in R8 and is autonomously required in Mil for the organization of columnar neurons.

**Fig. 4.**
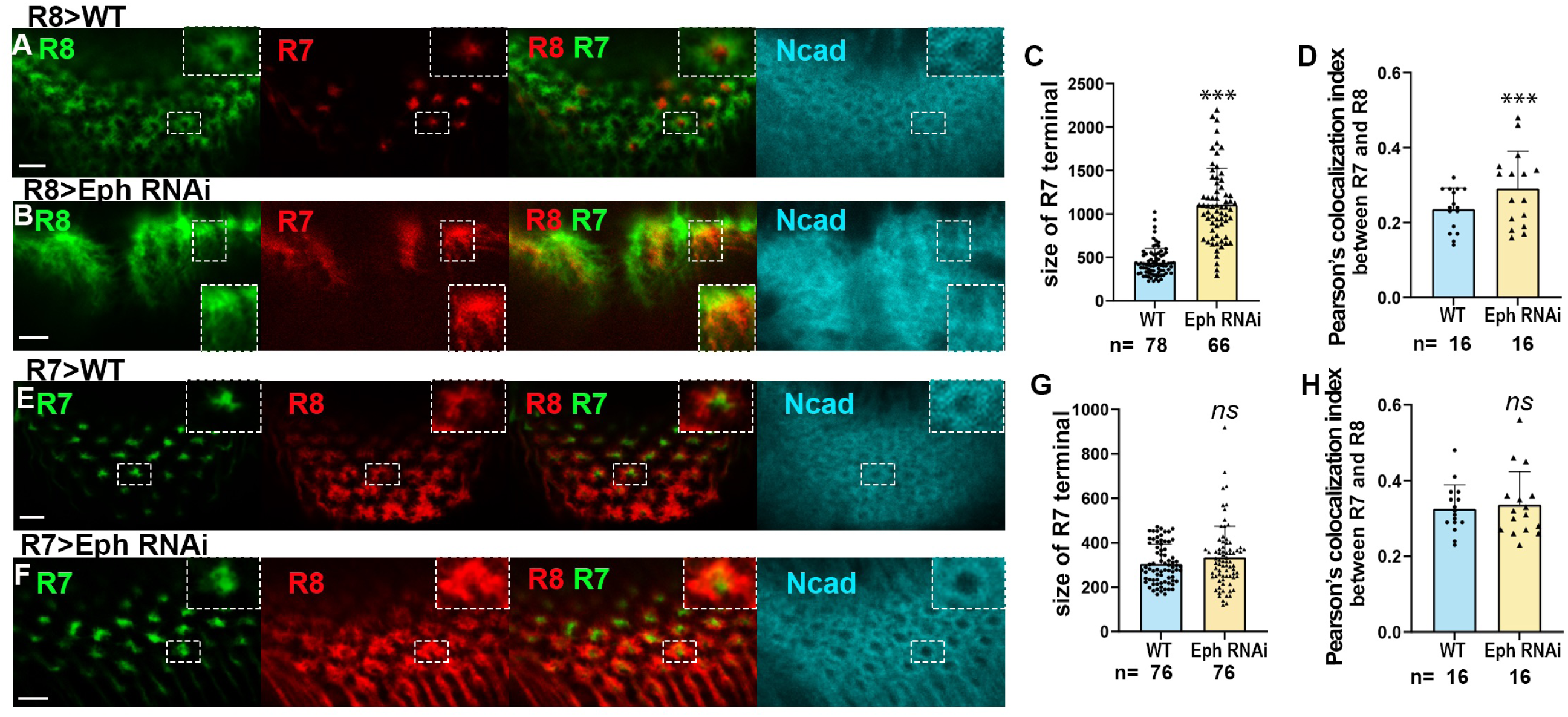
Eph expressed in R8 regulates the organization of columnar neurons. (**A-B**) *Eph* knock-down in R8 disturbs the separation of R7 (R7-GFP, green) and R8 (R8-RFP, red) terminals and column formation (Ncad, blue). (**C**) Quantification of the size of R7 terminals (in **A, B**). (**D**) Colocalization of the terminals of R7 and R8 (in **A, B**). (**E, F**) *Eph* knock-down in R7 did not affect the separation of R7 and R8 terminals and column formation (Ncad, blue). (**G**) Quantification of the size of R7 terminals (in **E, F**). (**H**) Colocalization of the terminals of R7 and R8 (in **E, F**). The dotted boxes are enlarged in the right upper panels (in **A, B, E, F**). Results were statistically analyzed using Welch’st-test. *** indicates p< 0.001, n.s. not significant. Scale bars, 5 μm.

Taken together, these results suggest that Ephrin/Eph forward signaling in the pairs of R7/R8 and R8/Mi1 and Eph/Ephrin reverse signaling in the pairs of R8/R7 are essential for column organization and that bi-directional signaling is activated between R7 and R8.

### The interaction between Ephrin and Eph is required for Ephrin phosphorylation

It is well-known that Eph/Ephrin reverse signaling is activated by the binding of Eph to Ephrin expressed in the neighboring cells. To test if Eph expressed in R8 is required for Ephrin phosphorylation in R7, clones expressing *Eph* RNAi were generated specifically in R8. Consequently, changes in Ephrin #1 signal were observed. Phosphorylated Ephrin was downregulated when *Eph* was knocked down in neighboring R8 terminals (Figs. 5A-C), suggesting that Eph/Ephrin interaction between R8 and R7 is necessary for Ephrin phosphorylation in R7.

**Fig. 5.**
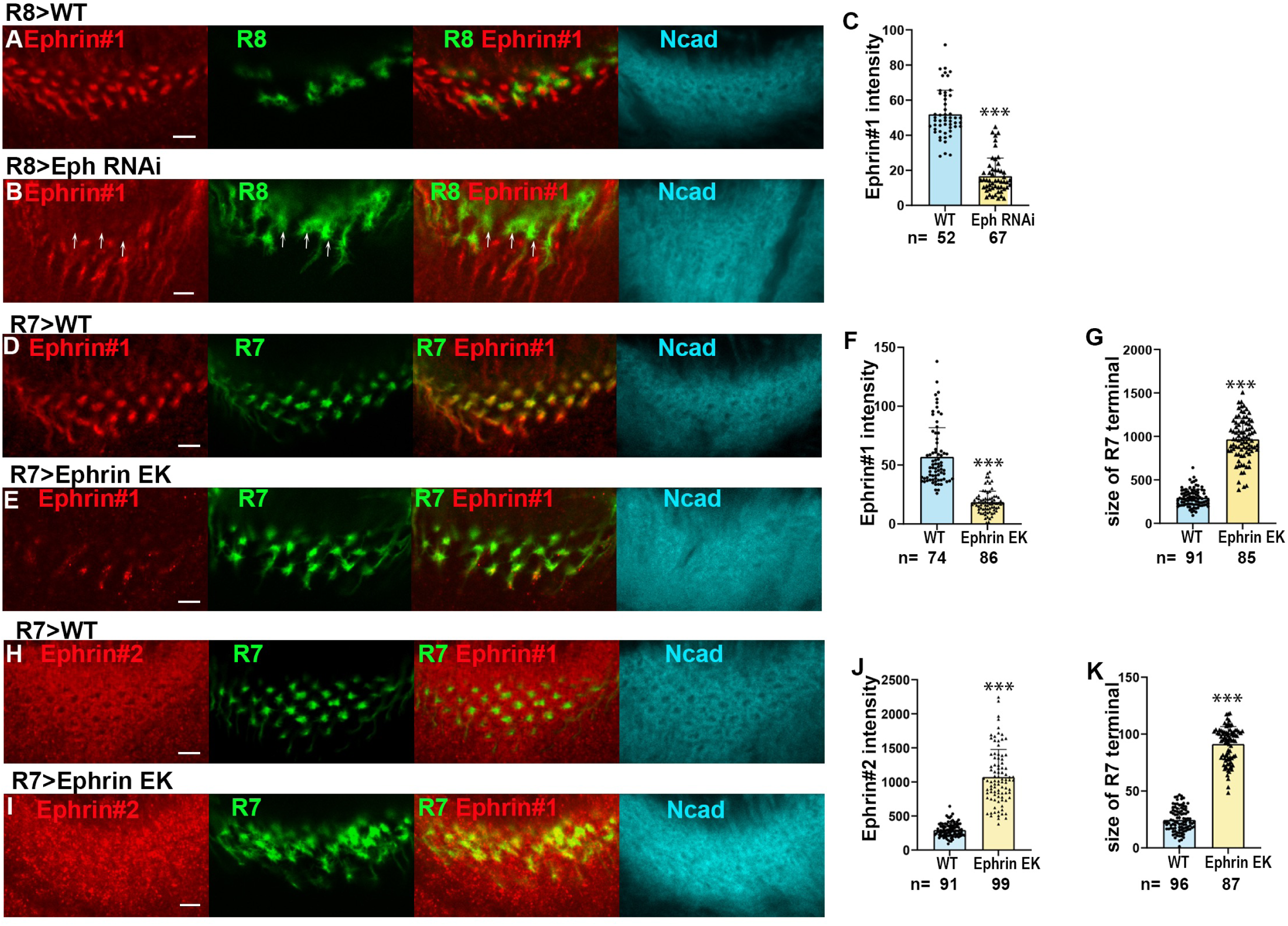
Interaction between Ephrin and Eph is required for Ephrin phosphorylation. (**A, B**) Ephrin#1 signals were downregulated nearby R8 terminals expressing *Eph* RNAi (white arrows). (**C**) Quantification of the signal intensities of Ephrin #1 (in **A, B**). (**D-K**) Expression of *Ephrin^EK^*in R7 caused down-regulation of Ephrin #1 signals (**E**), up-regulation of Ephrin #2 signals (**I**), and enlargement of R7 terminals (**E, I**). (**F, J)** Quantification of the signal intensities of Ephrin #1 (in **D, E**) and Ephrin #2 (in **H, I**) in R7 terminals. (**G, K**) Quantification of the size of R7 terminals (in **D, E, H, I**). All results were statistically analyzed using Welch’s t-test. *** indicates p< 0.001, n.s. not significant. Scale bars, 5 μm.

It has been reported that E320K mutation in the extracellular domain of Ephrin inhibits its binding to Eph in *trans* (*26*). Overexpression of *Ephrin^E320K^* (*Ephrin^EK^*) in R7 caused downregulation of Ephrin #1 signals and up-regulation of Ephrin #2 signals in R7 terminals, suggesting that the binding of Eph to Ephrin is required for Ephrin phosphorylation (Figs. 5D-K).

### Tyrosine phosphorylation is required for Ephrin reverse signaling

To test if Eph/Ephrin reverse signaling depends on tyrosine phosphorylation of Ephrin, we generated a mutant form of *Ephrin* in which the tyrosine residues in the cytoplasmic domain were mutated to phenylalanine (*Ephrin^2YF^*) (Fig. 6F). When Ephrin2YF was overexpressed in R7, Ephrin #1 signal was decreased and Ephrin #2 signal was increased (Figs. 6A-D). In addition, R7 terminals were expanded and disorganized, suggesting that tyrosine phosphorylation is required for Ephrin reverse signaling (Figs. 6A-I). However, the tyrosine residues may not be the only amino acids that are phosphorylated in the cytoplasmic domain of Ephrin, because Ephrin #1 signal was more significantly decreased when EphrinCD was expressed in R7 than when Ephrin2YF was expressed (Fig. S6).

**Fig. 6.**
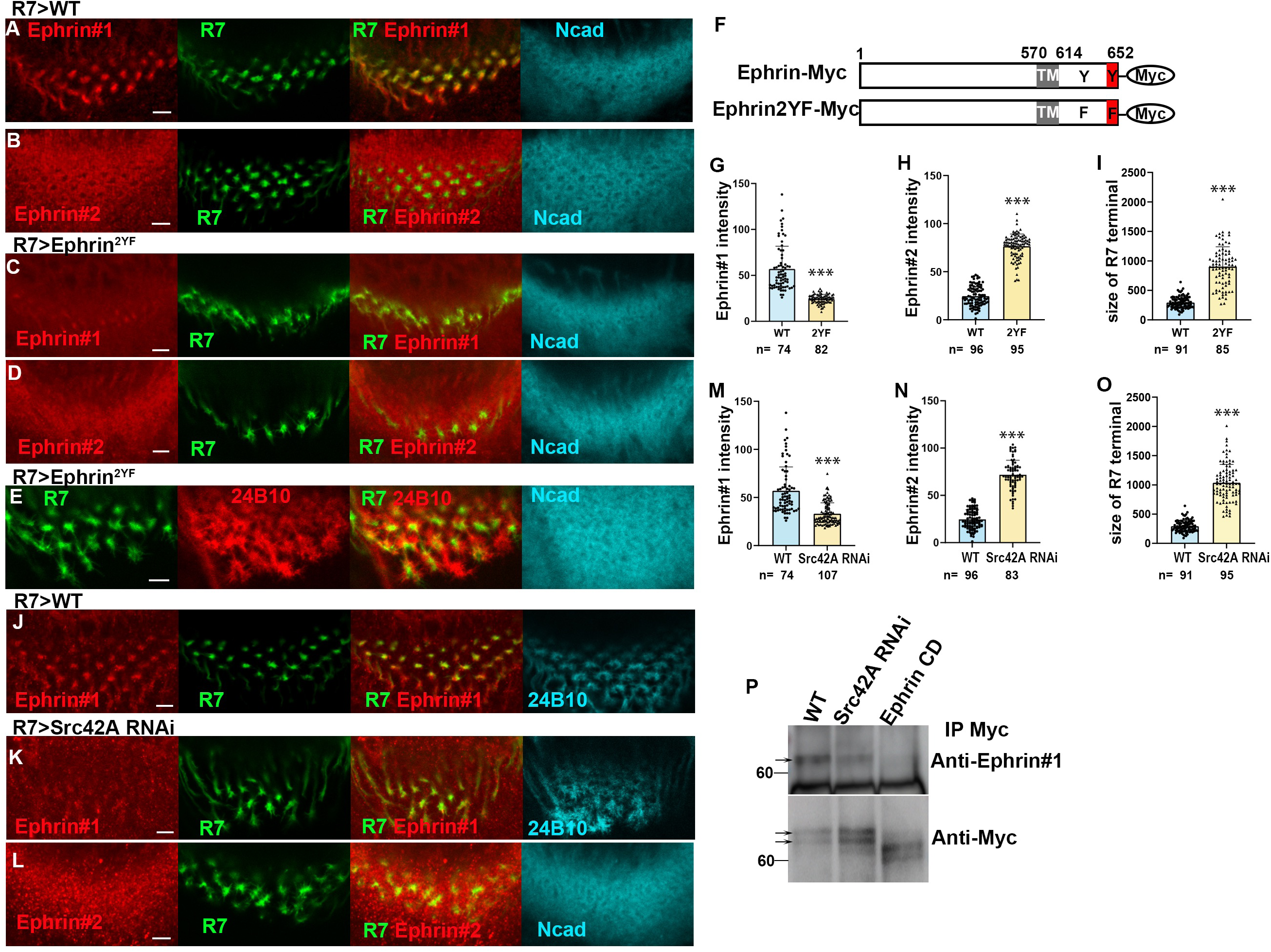
Tyrosine phosphorylation of Ephrin under the control of Src is required for Eph/Ephrin signaling. (**A-E**) Expression of *Ephrin^2YF^* in R7 decreased Ephrin #1 signals (**C**), increased Ephrin #2 signals (**D**), and enlarged the terminals of R7 (**B, D, E**). (**F**) Two tyrosine residues in the cytoplasmic domain of Ephrin were mutated to phenylalanine in *Ephrin^2YF^*. (**G-I**) Quantification of the signal intensities of Ephrin #1 (in **A, C**), Ephrin #2 (in **B, D**), and the size of R7 terminals (in **A-E**). (**J-L**) *Src42A* knocked-down in R7 decreased Ephrin #1 signals (**K**) and increased Ephrin #2 signals (**L**). 24B10, blue (**J, K**). Ncad, blue (**L**). (**M-O)** Quantification of the signal intensities of Ephrin#1 (in **J, K**), Ephrin #2 (in **B, L**), and the terminals of R7 (in **J, K**). Results were statistically analyzed using Welch’s t-test. *** indicates p< 0.001, n.s. not significant. Scale bars, 5 μm. (**P**) Ephrin#1 signals were decreased upon *Src42A RNAi* and *EphrinCD* expression after immunoprecipitation.

### Src kinases are required for Ephrin phosphorylation

To identify tyrosine kinases that phosphorylate Ephrin, we performed R7 specific RNAi screening focusing on twelve genes encoding non-receptor tyrosine kinases. We found that knock-down of *Src42A*, one of the two *Drosophila* Src homologues, decreased Ephrin #1 signals and increased Ephrin #2 signals in the terminals of R7 axons (Figs. 6J-O). The decrease in phosphorylated Ephrin was confirmed by Western blotting followed by immunoprecipitation from larval brain extracts (Fig. 6P). These results are consistent with the observation that Src positively regulates Ephrin-B phosphorylation and mediates Eph/Ephrin-B reverse signaling *in vitro* (*20, 27*).

In *Drosophila*, Src42A and Src64B play redundant roles in multiple aspects of development (*28–30*). However, Ephrin #1 signal was not significantly downregulated upon *Src64B* knock-down in R7. Since downregulation of Ephrin #1 signal was enhanced upon double knock-down of *Src42A* and *Src64B* (Fig. S7), both are required for Ephrin phosphorylation, while Src42A plays more critical roles compared to Src64B.

### Fas2 is required for Ephrin phosphorylation and Eph/Ephrin reverse signaling

Fasciclin II (Fas2) is a member of the immunoglobulin superfamily and is homologous in structure and function to neural cell adhesion molecule, NCAM, a pivotal regulator of axon growth, fasciculation, and cell adhesion (*31*). It was reported that NCAM cooperates with the EphrinA/EphA system in restricting arborization of GABAergic interneurons in mouse prefrontal cortex (*32*). However, it is not known if NCAM and/or Fas2 is involved in Eph/Ephrin reverse signaling. Interestingly, we found that Fas2 is colocalized with Ephrin #1 signals in R7 (Fig. 7A). When *Fas2* was knocked down in R7, Ephrin #1 signal was downregulated and Ephrin #2 signal was upregulated, indicating that Fas2 is required for Ephrin phosphorylation (Figs. 7B-H). The Fas2 dependence of Ephrin phosphorylation was confirmed by immunoprecipitation of Ephrin-Myc expressed under the control of *GMR-Gal4* followed by Western blotting (Fig. 7L). In addition, the R7 terminals showed irregular morphology and larger size in the *Fas2* RNAi condition (Figs. 7I-K). The column morphology was also disrupted as visualized by Ncad (Figs. 7E, F, I, J). Taken together, these data suggest that Fas2 is required for the Ephrin phosphorylation and Eph/Ephrin reverse signaling in R7.

**Fig 7.**
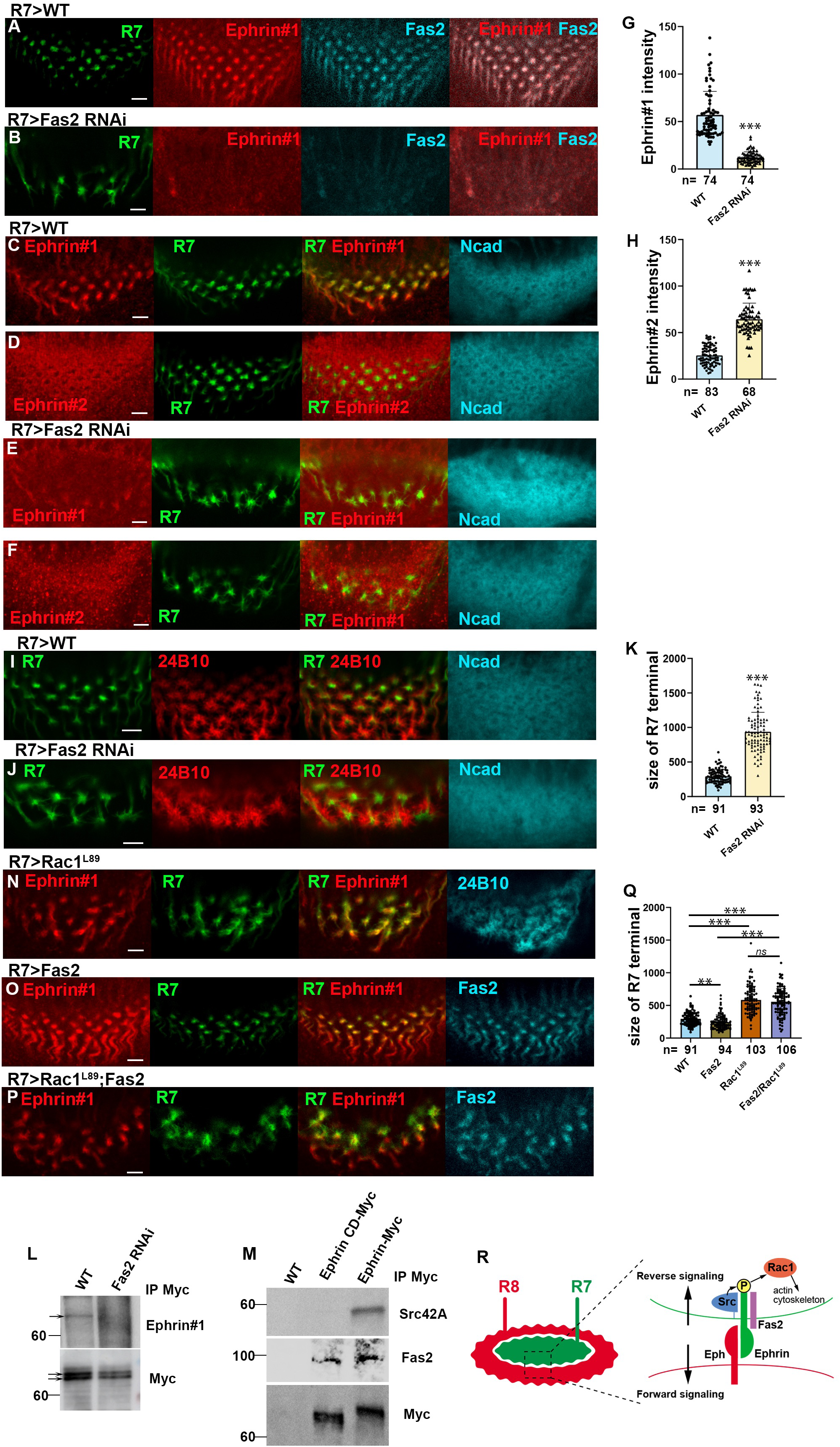
Fas2 is required for Ephrin phosphorylation in R7. (**A, B**) Downregulation of Ephrin #1 signals (red) and Fas2 (blue) upon *Fas2 RNA*i in R7. (**C-F**) Down-regulation of Ephrin#1 (red in **C, E**) and up-regulation of Ephrin#2 signals (red in **D, F**) upon *Fas2 RNAi* in R7. Ncad, blue. (**G-K**) Quantification of the intensities of Ephrin #1 (in **C, E**) and Ephrin #2 (in **D, F**). (**I, J)** The terminals of R7 (green) and R8 (24B10, red) were disorganized upon *Fas2 RNAi* in R7. (**K**) Quantification of the size of R7 terminals (in **I, J**). (**L**) Ephrin#1 signals decreased upon *Fas2 RNAi* after immunoprecipitation (arrows). (**M**) Src42A and Fas2 were co-immunoprecipitated by Ephrin-Myc, while only Fas2 was co-immunoprecipitated by EphrinCD-Myc. (**N**) The terminals of R7 (green) and R8 (24B10, blue) were disorganized upon *Rac1^L89^*expression in R7. (**O**) Ectopic expression of *Fas2* (blue) in R7 reduced the size of R7 terminals. (**P**) Ectopic expression of *Fas2* (blue) together with *Rac1^L89^* enlarged R7 terminals. (**Q**) Quantification of the size of R7 terminals (in **I**, **N-P**). Results were statistically analyzed using Welch’s t-test. *** indicates p< 0.001, n.s. not significant. Scale bars, 5 μm. (**R**) Schematic drawing of Ephrin phosphorylation in R7 depending on the complex of Src, Fas2 and Ephrin triggered by Eph expressed in R8.

Next, to verify if Fas2 and Src42A interact with Ephrin, we performed a coimmunoprecipitation experiment *in vivo*. When Ephrin-Myc was immunoprecipitated with Myc antibody, coimmunoprecipitation of endogenous Fas2 and Src42A was detected by Western blotting (Fig. 7M). These results indicate that Src42A and Fas2 interact with Ephrin, which leads to Ephrin phosphorylation followed by Eph/Ephrin reverse signaling.

### Rac1 mediates Eph/Ephrin reverse signaling to control the organization of columnar neurons

Previous studies demonstrated that Ephrin-B reverse signaling plays a role in modulating axon retraction and pruning through a small GTPase, Rac1/3 (*21, 33, 34*). To investigate the role of Rac1 in R7 terminal organization, we expressed a dominant-negative form of Rac1 (Rac1^L89^) in R7, which resulted in the disorganization of the terminals of R7 and R8, mimicking the effects observed when Ephrin or Fas2 function was impaired in R7 (Figs. 3B, 7J, 7N). Notably, ectopic expression of Fas2 in R7 led to a reduction in the size of R7 terminals compared to the control (Figs. 7C, 7O, 7Q), suggesting that Fas2 promotes Eph/Ephrin reverse signaling, which in turn reduces the size of R7 terminals.

To examine whether Rac1 acts as an essential effector downstream of Eph/Ephrin reverse signaling, we overexpressed Fas2 together with the dominant negative form of Rac1 in R7. Since the reduced R7 terminals caused by Fas2-overexpression were suppressed by co-expression of Rac1 dominant negative, the results suggest that Rac1 acts downstream of Eph/Ephrin reverse signaling induced by Fas2 (Figs. 7O, 7P, 7Q). Thus, Rac1 acts as a downstream effector of Eph/Ephrin reverse signaling to regulate the organization of columnar neurons.

## Discussion

In this study, we demonstrated that the bi-directional repulsive Ephrin/Eph signal shapes the columnar unit by organizing the morphology and segregation of columnar neurons in the fly brain. Furthermore, we found that the binding of Fas2 and Src kinase to Ephrin triggers tyrosine phosphorylation of Ephrin, which initiates Eph/Ephrin reverse signaling through Rac1. We presented a unified picture of the complex molecular interactions in the defined context of column formation for the first time.

During cerebral cortex development, different types of excitatory neurons originating from the proliferative ventricular zone radially migrate to form a cellular infrastructure of columns (*35, 36*). Ephrin/Eph signaling was shown to be essential for columnar distribution and the proper assembly of cortical neurons (*21, 22*). Similarly, in the fly brain, R7 and R8 axons project from the retina to the medulla where their terminals comprise the medulla columns together with other axons and dendrites of columnar neurons such as Mi1. Our study proved that phosphorylated and unphosphorylated Ephrin show distinct distribution patterns in columnar neurons. Ephrin in R7 was phosphorylated on the tyrosine residues under the control of Eph expressed in adjacent R8 to trigger Eph/Ephrin reverse signaling. Forward signaling of Eph in R8 and reverse signaling of Ephrin in R7 control the morphogenesis of the terminals of core columnar neurons (Fig. 7N). Unphosphorylated Ephrin in R8 regulates forward signaling of Eph in Mi1 to control columnar neuron organization as well (Figs. S4, S5).

In vertebrates, transmembrane Ephrin-B acts as a receptor, in part mediated by its tyrosine phosphorylation. Ephrin-B has highly conserved tyrosine residues that are phosphorylated upon interaction with Eph (*37*). *Drosophila* Ephrin has a sequence homologous to Ephrin-B (*19, 23*). Tyrosine and Serine phosphorylation of Ephrin-B has been shown to function in two independent events regulating different aspects of Ephrin-B reverse signaling (*38*). In our study, tyrosine phosphorylation was shown to be required to regulate Ephrin reverse signaling by using a mutant form of Ephrin in which the two tyrosine residues in the cytoplasmic domain are mutated (Fig. 6).

We identified Src42A, one of the *Drosophila* Src family kinases, as a candidate that acts as a tyrosine kinase of Ephrin (Figs. 6J-P). Ephrin phosphorylation and its reverse signaling was compromised by knocking down *Src42A* in R7. Consistent with this, Src family kinases (SFKs) were shown to positively regulate Ephrin-B phosphorylation and its phosphotyrosine-mediated reverse signaling in vertebrates (*20*). SFKs may be recruited to Ephrin-B in an Eph dependent manner. We demonstrated the binding of Src42A to the cytoplasmic domain of Ephrin (Fig. 6P). Thus, the binding of Eph expressed in R8 to Ephrin expressed in R7 may lead to the recruitment of Src42A, which may induce the downstream event of Eph/Ephrin reverse signaling. Since the expression of Src42A is uniform in the columnar neurons, we propose that there is an additional regulatory factor specifically expressed in R7 that controls the interaction between Src42A and Ephrin.

Interestingly, Fas2 is colocalized with phosphorylated Ephrin in R7. We demonstrated that Fas2 binds to the extracellular domain of Ephrin and is required for Ephrin phosphorylation (Fig. 7). Thus, Ephrin binds both Src42A and Fas2 in R7 resulting in Ephrin phosphorylation and activation of Ephrin reverse signaling triggered by Eph expressed in R8. Fas2 may play a critical role in the formation of the complex of Ephrin/Src42A/Fas2 in the presence of Eph in adjacent cells. A previous study showed that NCAM, a homologue of Fas2, forms a complex with EphA3, which is necessary for EphrinA5/EphA3 signaling and plays a crucial role in restricting arborization of GABAergic interneurons in the mouse prefrontal cortex (*32*). However, its involvement in Eph/Ephrin reverse signaling was not known. Based on our data, Fas2 probably initiates the recruitment of Src42A to the Ephrin cytoplasmic domain under the control of Eph. In vertebrates, the PDZ-binding motifs in the cytoplasmic tail of Ephrin-B are involved in its tyrosine phosphorylation (*39*). However, the cytoplasmic tail of *Drosophila* Ephrin does not contain the PDZ binding sequence (*23*), suggesting that an alternative mechanism may induce Ephrin activation in *Drosophila*. Despite these differences, the tertiary complex of Ephrin/Src/NCAM (Fas2), or the quaternary complex including Eph, may play important roles in stabilizing Eph/Ephrin reverse signaling during column formation.

We next addressed a downstream effector of Eph/Ephrin reverse signaling that could regulate cytoskeletal organization at the terminals of columnar neurons. The small GTPase, Rac1, was implicated in cell migration and neuronal morphogenesis through cytoskeletal organization (*40–42*), and was shown to be involved in Ephrin-B reverse signaling (*33*). We demonstrated that the effect of Fas2 overexpression in R7 to reduce the size of R7 terminals was suppressed by co-expression of Rac1 dominant negative (Fig. 7), suggesting that Rac1 acts as a downstream effector of Eph/Ephrin reverse signaling to control the organization of columnar neurons.

Thus, we presented a unified picture of the complex molecular interactions in the defined context of column formation. It is very likely that the molecular mechanisms we have demonstrated in a series of *in vivo* experiments are evolutionarily conserved from flies to mammals. Our findings open up a new avenue of research that will help to broaden our understanding of the mechanisms of column formation and other related developmental processes.

## Materials and Methods

### Fly Strains

Standard *Drosophila* medium was used to maintain fly strains at 25°C. Both male and female flies were used for all experiments. The following fly strains were used: *UAS-Ephrin RNAi*, *UAS-Ephrin^E320K^*, *UAS-Eph-Myc*, *UAS-Eph RNAi* (from Dr. Takahiro Chihara), *UAS-Src42A RNAi* (BDSC#44039), *UAS-Src64B RNAi* (BDSC#51772), *UAS-Fas2 RNAi* (BDSC#34084), *UAS-Rac1^L89^* (BDSC#6290). *bshM-Gal4 UAS-myrGFP* (Mi1-GFP), *bshM-LexA LexAop-myrTomato* (Mi1-RFP), *sevEnS-Gal4 UAS-myrGFP* (R7-GFP), *sevEnS-LexA LexAop-myrTomato* (R7-RFP), *sensF2-Gal4 UAS-myrGFP* (R8-GFP), and *sensF2-LexA LexAop-myrTomato* (R8-RFP) were generated in our previous work^3^. *sens-FLPase; GMR-FsF-Gal4 UAS-myrGFP* was used to perform R8 specific knockdown (*43*).

### Generation of *UAS-EphrinCD-Myc* and *UAS-Ephrin^2YF^-Myc* vectors and strains

The *Ephrin-Myc* plasmid was obtained as a gift from Dr. Takahiro Chihara. *Ephrin-Myc* cDNA was amplified using forward primer *F1* and reverse primer *R1* and then inserted into the *NotI* and *KpnI* sites of the *pBSKS* vector. To create a cytoplasmic deletion of *Ephrin*, we used the *pBSK-Ephrin* vector as a template and carried out PCR amplification using primer pair *CD_Fwd* and *CD_Rev.* The resulting PCR product was then self-ligated using NEBuilder HiFi DNA Assembly Reaction (NEB) following the standard protocol. Similarly, we generated two-point mutations *Y650F* and *Y612F* of *Ephrin* using primer pairs *2YT_Fwd1/2YT_Rev1* and *2YT_Fwd2/2YT_Rev2*, respectively. The resulting PCR products were treated with *DpnI* (NEB) to remove the template plasmid DNA and then self-ligated using Ligation High Ver2 (Takara). All PCR products and enzyme-digested vectors were purified using QIAquick PCR Purification Kit (Qiagen), and the resulting vectors were transformed into ECOSTM Competent *E. coli DH5*α (Nippon Gene). After confirming the sequences, the cytoplasmic deletion of *Ephrin* and the two-point mutated *Ephrin* segments were excised using *NotI* and *KpnI* and re-inserted into the *pUAST* vector. The resulting P-element vectors were then microinjected by GenetiVision, USA.

*F1*: aattggagctccacc**<u>gcggccgc</u>**ATTATGCAAGAACGATCAAAGCA

*R1*: gggaacaaaagctg**<u>ggtacc</u>**CTAGACTAGTGGATCCCCCGG

*CD_Fwd*: attcgctctagacggTTCCATCGATTTAAAGCTATGGAGC

*CD_Rev*: tccgtctagagcgaatAAGATAGTGAATGCCAAGAATTGC

*2YT_Fwd1*: **T**TGACCGGTTCCATCGATTTAAAGC

*2YT_Rev1:* **A**ATTCAATAGTGCCAGCATTC

*2YT_Fwd2:* **T**TAGTCCTGGAATGGTTGAA

*2YT_Rev2:* **A**ATCGCTGCACCGTGGTTTGC

### Histochemistry

Immunohistochemistry was performed as described previously ^1^. Details are available upon request. Brains were dissected in PBS, transferred to an ice-cold 0.8% formaldehyde/PBS solution, and fixed in 4% formaldehyde/PBS at room temperature for 30-60 minutes. The brains were washed in PBT (0.3% TritonX in PBS) and blocked in a 5-10% donkey normal serum/PBT solution at room temperature for 30-60 minutes. The primary antibody reaction was performed in a solution containing primary antibodies and 1% normal serum in PBT at 4°C overnight. The brains were washed in PBT. The secondary antibody reaction was performed in a solution containing secondary antibodies and 1% normal serum in PBT at 4°C overnight. After washing in PBT and PBS, the brains were mounted in VECTASHIELD (Vector Laboratories).

The following primary antibodies were used: rat anti-Ncad (1:20, Developmental Study Hybridoma Bank), mouse anti-Chaoptin (24B10, 1:20, Developmental Study Hybridoma Bank), mouse anti-Fas2 (1:20; Developmental Study Hybridoma Bank), rabbit anti-Myc (1:1000, Santa Cruz Biotechnology), and rabbit anti-Src42A (1:200; from Dr. Tetsuya Kojima). Ephrin antibodies were generated in rabbits against the synthetic peptide, CNGMFDQNAGTIEYDR, which corresponds to the intracellular domain of Ephrin (WAKO, Japan; Fig. 2F).

The following secondary antibodies were used: anti-mouse Cy3 (1:200; Jackson ImmunoResearch 715-165-151), anti-mouse Cy5 (1:200; Jackson ImmunoResearch 715-175-151), anti-rat Alexa647 (1:100; Jackson ImmunoResearch 712-605-150), and anti-rabbit Alexa546 (1:200; Invitrogen A-11035). Confocal images were obtained by Zeiss LSM880 and processed using ZEN, Fiji, and Adobe Photoshop.

### Coimmunoprecipitation and Immunoprecipitation

Dissected brain samples were collected in PBS and lysed with 1x lysis buffer containing 1× protease inhibitor cocktail (10x lysis buffer, Nacalai Tesque, Japan) and with or without phosphatase inhibitor (Nacalai tesque) in the presence of 1 mM PMSF and 10 mM NaF. Anti-Myc-tag mouse antibody Magnetic beads were added and immunoprecipitated according to the manufacturer’s protocol (MBL and ThermoFisher). Western blotting was performed according to standard techniques.

### Statistics and reproducibility

Distinct brain areas or samples were measured and analyzed as indicated below for quantification and statistical analysis. A two-sided t-test was used for the statistical test. Image intensities were not artificially processed, except as otherwise noted. At least six columns from more than ten brain samples were examined. When statistics were not applicable, experiments were independently repeated at least three times with similar results.

### Image quantification

Signal intensity was quantified within the indicated rectangle areas using Fiji. The Coloc 2 function of Fiji was used to calculate the Pearson’s *R* value to quantify the colocalization between two different signals. The graphs of quantified data were generated by Prism.

## Acknowledgements

We thank Takahiro Chihara for fly strains and vectors, and Tetsuya Kojima for antibody. We thank Takahiro Chihara, Steven R. Davis, and Atsushi Sugie for critical comments on the manuscript. We thank members of Sato lab for their support and helpful discussion. We thank the Bloomington Stock Center, Vienna Drosophila RNAi Center, and DGRC Kyoto, for fly strains and the Developmental Study Hybridoma Bank (DSHB) for antibodies. This work was supported by Grant-in-Aid for Scientific Research (B), Grant-in-Aid for Transformative Research Areas (A), Early-Career Scientists and JSPS fellow from MEXT (20H01823, 21H02484, 22H05169 and 22H05621 to M.S., and 22K15121 and 22F32073 to M.W.).

## Author contributions

M.W. and M.S. conceived and designed the experiments. M.W., X.H., Y.L., R.T. and M.S. performed the experiments. M.W., X.H., and M.S. acquired, analyzed, and interpreted the data. M.W. and M.S. wrote the manuscript.

---

**Fig. S1.**
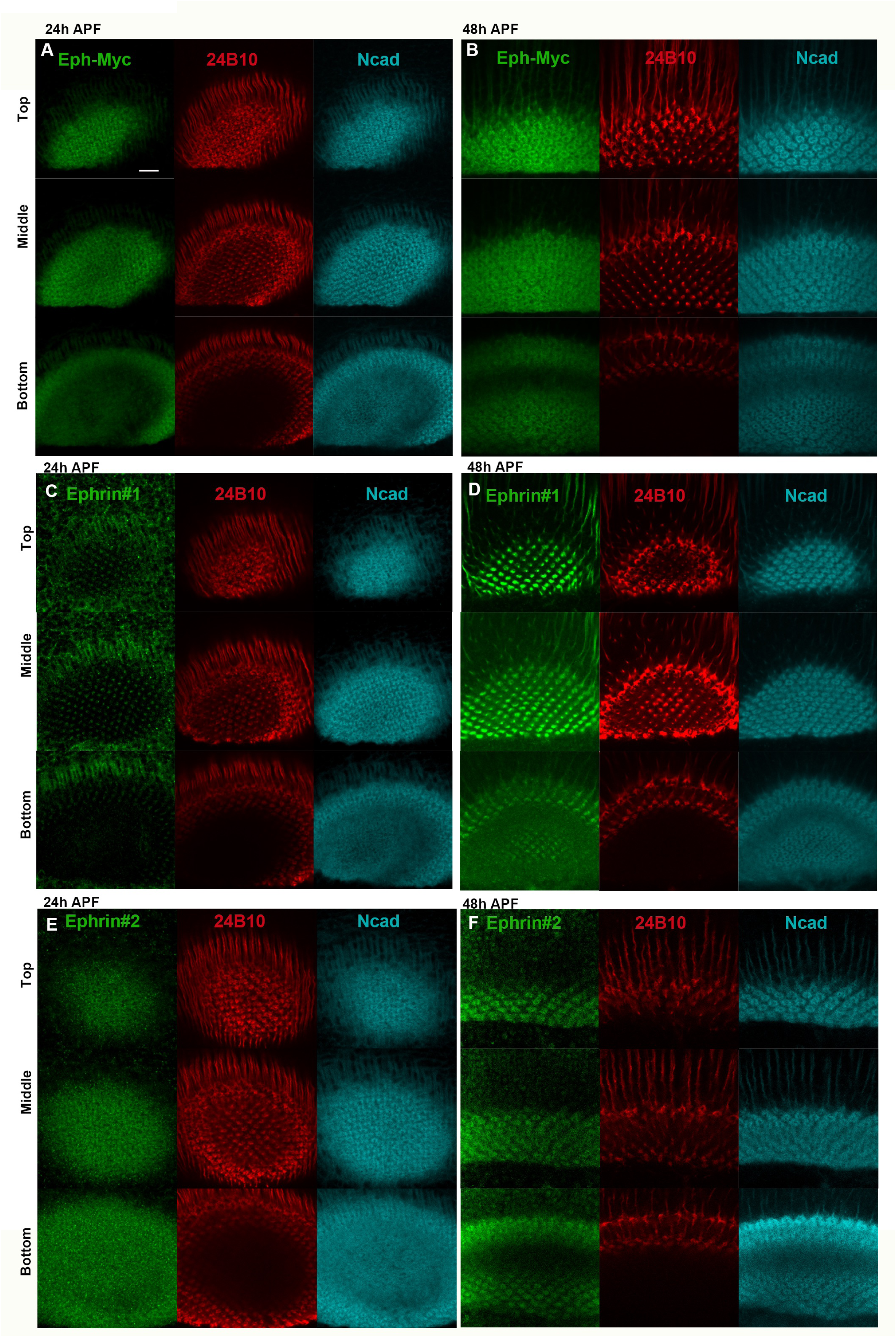
Eph and Ephrin distributions during pupal development. (**A, B**) Eph-Myc shows a donut-like pattern (Myc, green) overlapping with Ncad (blue) at 24 h (**A**) and 48 h APF (**B**). (**C, D)** Ephrin #1 shows a dot-like pattern inside the donut-like domain of Ncad (blue) at 24 h (**C**) and 48 h APF (**D**). (**E, F**) Ephrin #2 shows a donut-like pattern overlapping with Ncad (blue) at 24 h and 48 h APF. 24B10 visualizes R7 and R8 terminals (red). Scale bars, 5 μm.

**Fig. S2.**
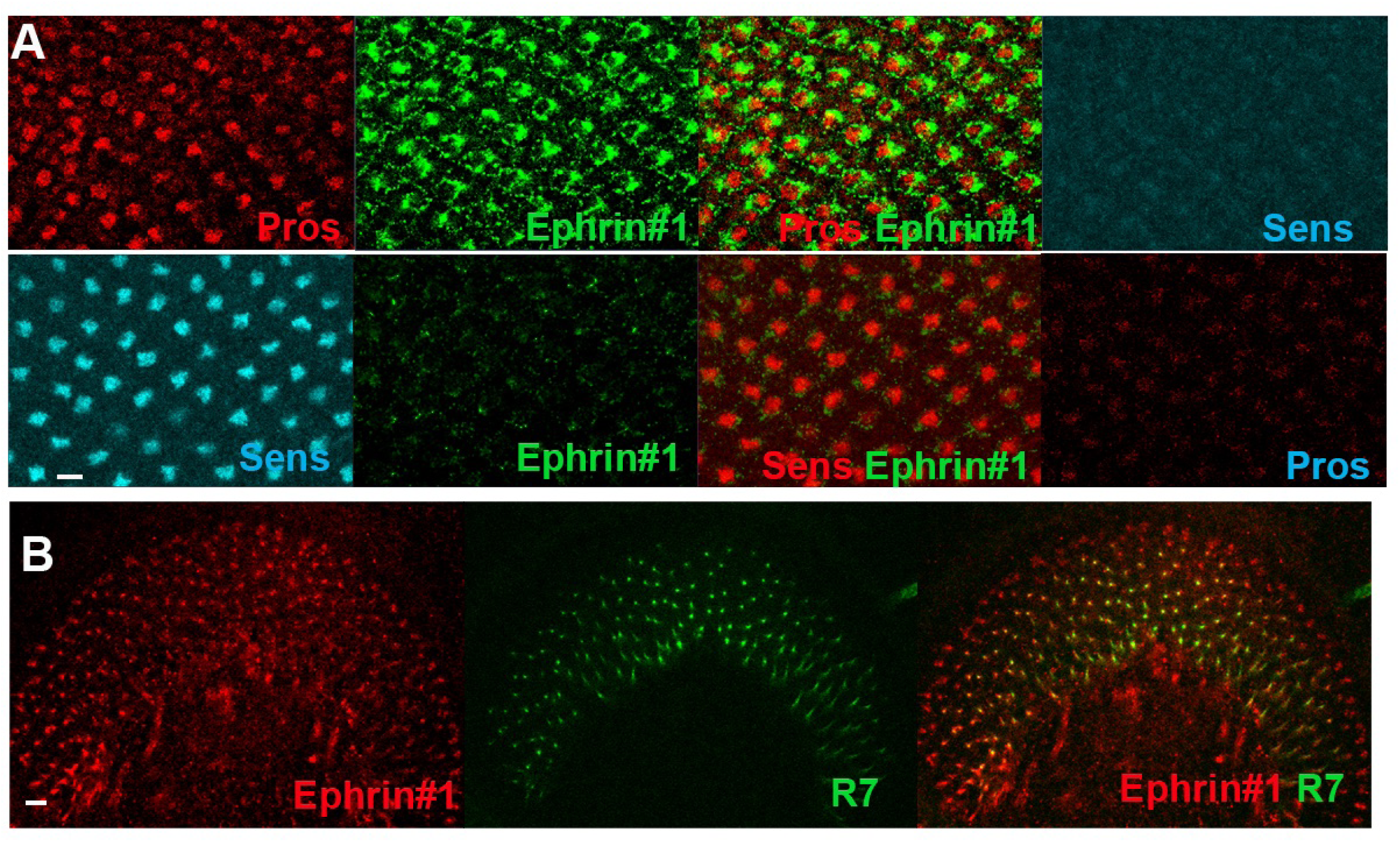
R7 is the source of Ephrin #1 signals. (**A**) In the late third larval instar eye disc, strong Ephrin #1 signals are found in R7 cell bodies visualized by Pros (red), but not in R8 visualized by Sens (blue). (**B**) In the late third larval instar lamina, Ephrin #1 signals (red) overlap with R7 axons (green). Scale bars, 5 μm.

**Fig. S3.**
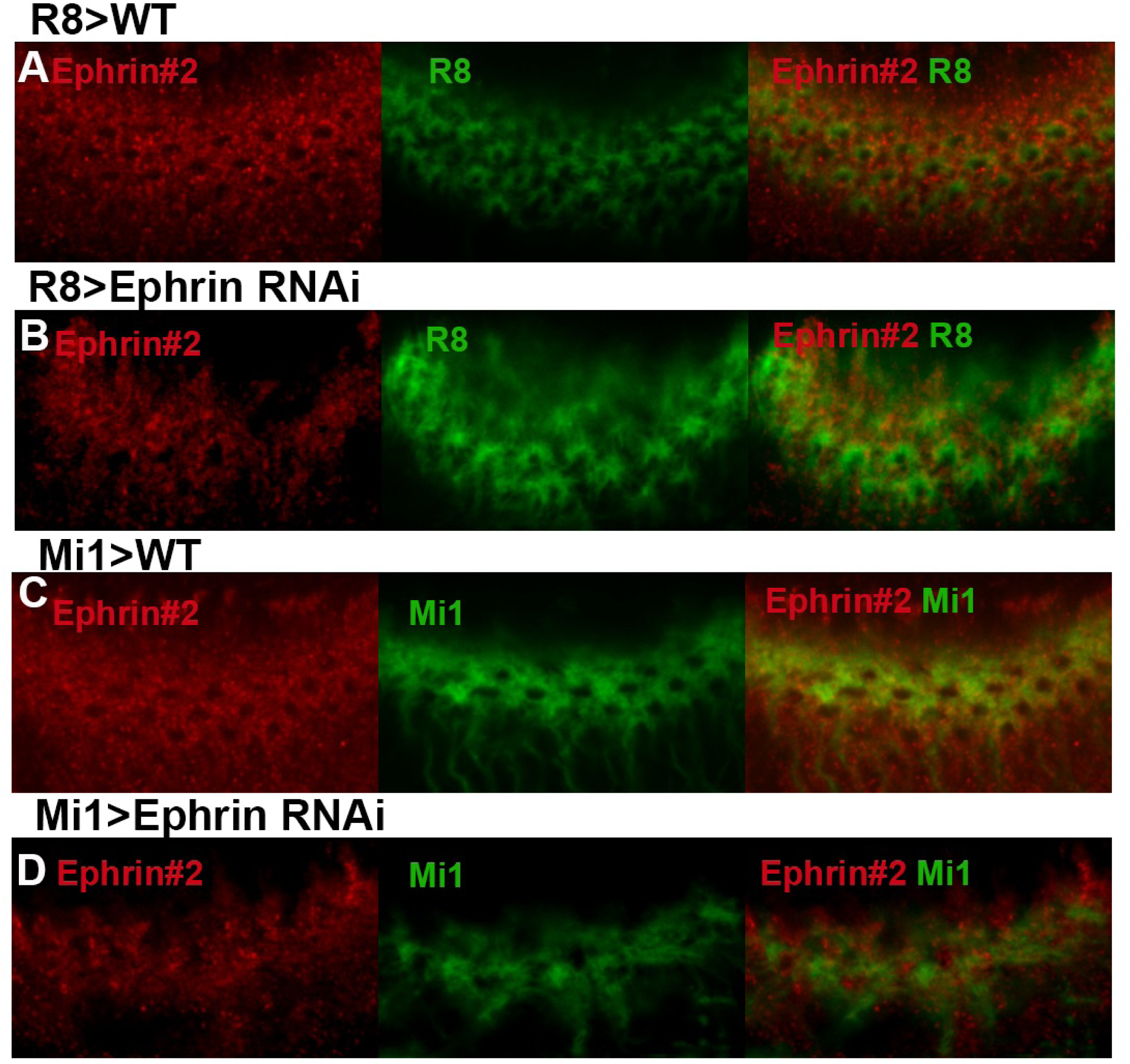
Ephrin#2 signals overlap with R8 and Mi1 terminals. (**A-D**) *Ephrin* knock-down in R8 (**B**) and Mi1 (**D**) reduced Ephrin #2 signals (red) in R8 and Mi1 terminals (R8-GFP and Mi1-GFP, green), respectively. Scale bars, 5 μm.

**Fig. S4.**
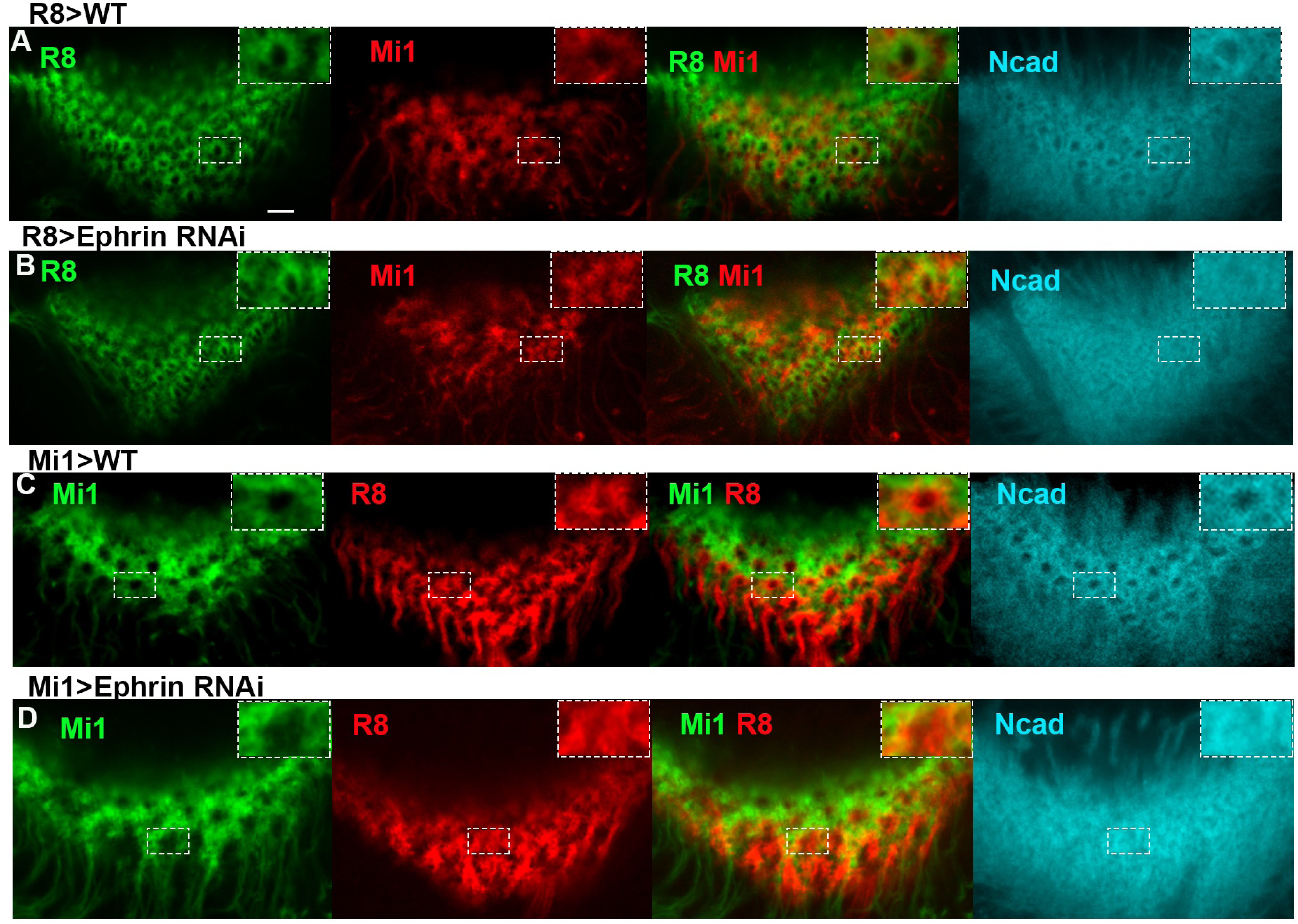
Ephrin expressed in R8 and Mi1 is non-autonomously required for column organization. (**A, B**) R8 and Mi1 terminals (R8-GFP, green and Mi1-RFP, red) were disorganized upon *Ephrin* knock-down in R8. (**C, D**) R8 and Mi1 terminals (R8-GFP, red and Mi1-RFP, green) were disorganized upon *Ephrin* knock-down in Mi1. Scale bars, 5 μm.

**Fig. S5.**
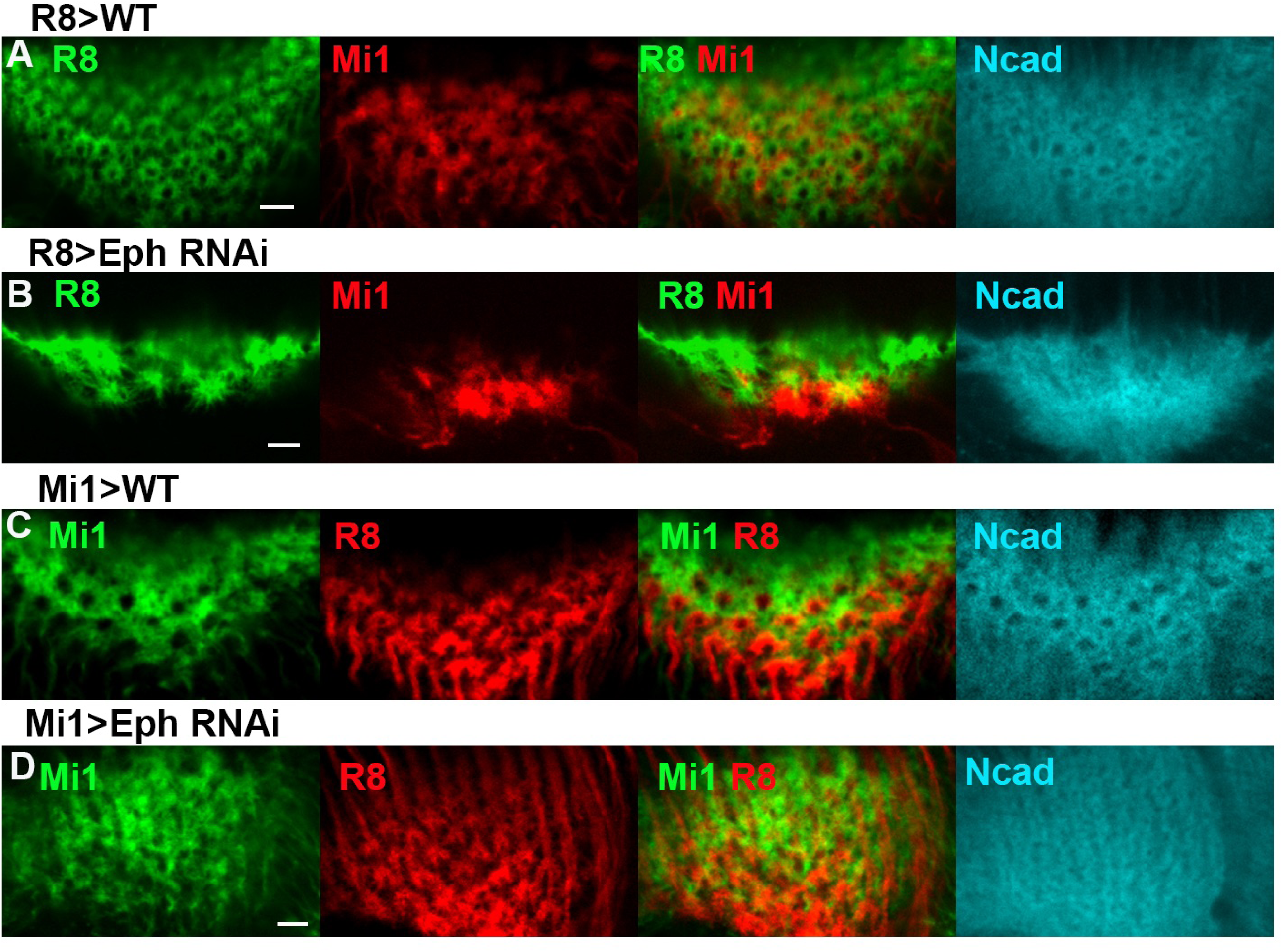
Eph expressed in R8 and Mi1 is required for column organization. (**A, B**) R8 and Mi1 terminals (R8-GFP, green and Mi1-RFP, red) were disorganized upon *Eph* knock-down in R8. (**C, D**) R8 and Mi1 terminals (R8-GFP, red and Mi1-RFP, green) were disorganized upon *Eph* knock-down in Mi1. Scale bars, 5 μm.

**Fig S6.**
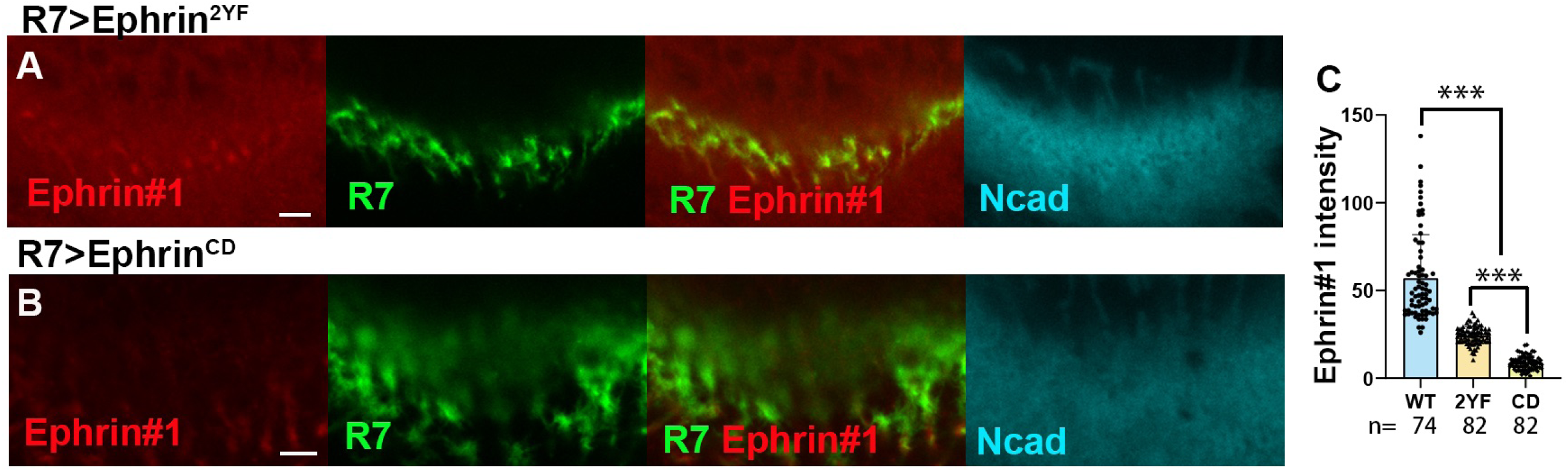
Expression of mutant forms of *Ephrin* suppresses Ephrin phosphorylation. (**A, B**) Ectopic expression of *Ephrin^2YF^* (**A**) and *Ephrin^CD^* (**B**) in R7 (R7-GFP, green) reduces Ephrin #1 signals (red). See Figure 2I as a control. (**C**) Quantification of Ephrin #1 intensity. *** indicates p< 0.001, n.s. not significant. Scale bars, 5 μm.

**Fig. S7.**
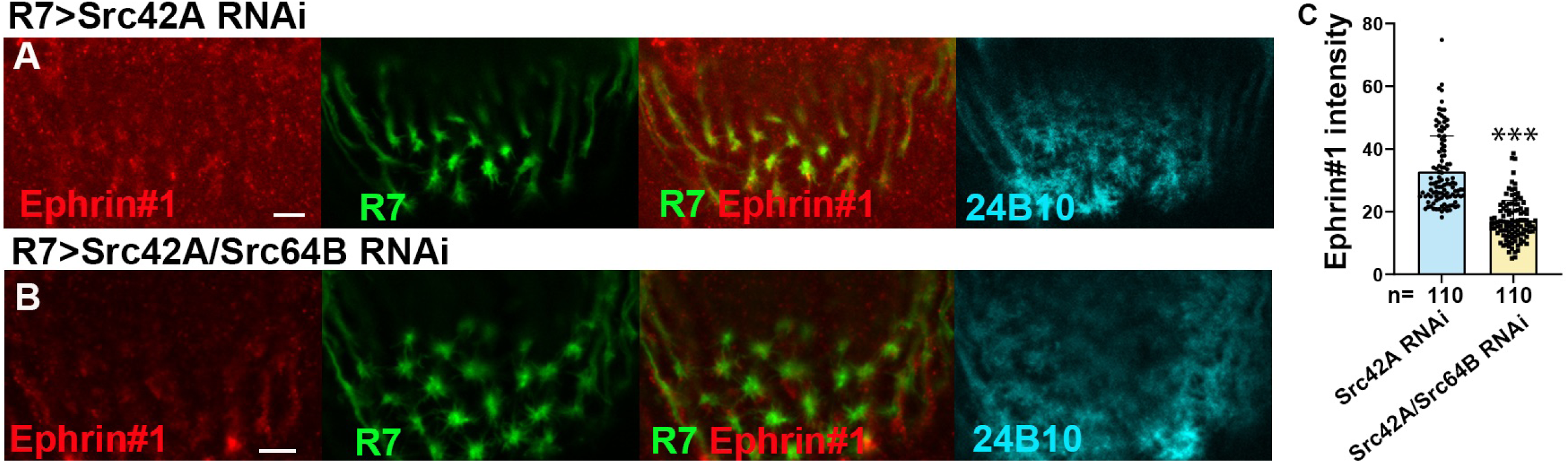
Src42A/Src64B double knockdown decreases Ephrin phosphorylation. (**A**) *Src42A* knock-down in R7 (R7-GFP, green) reduced Ephrin #1 signals. See Figure. 2I as a control. (**B**) *Src42A* and *Src64B* double knock-down in R7 (R7-GFP, green) further reduced Ephrin #1 signals. (**C**) Quantification of Ephrin #1 intensity. *** indicates p< 0.001, n.s. not significant. Scale bars, 5 μm.

