## Supplemental Figures for "Src/Fas2-dependent Ephrin phosphorylation initiates Eph/Ephrin reverse signaling through Rac1 to shape columnar units in the fly brain"

### Supplementary Materials

#### Supplementary Figures

Figure S1

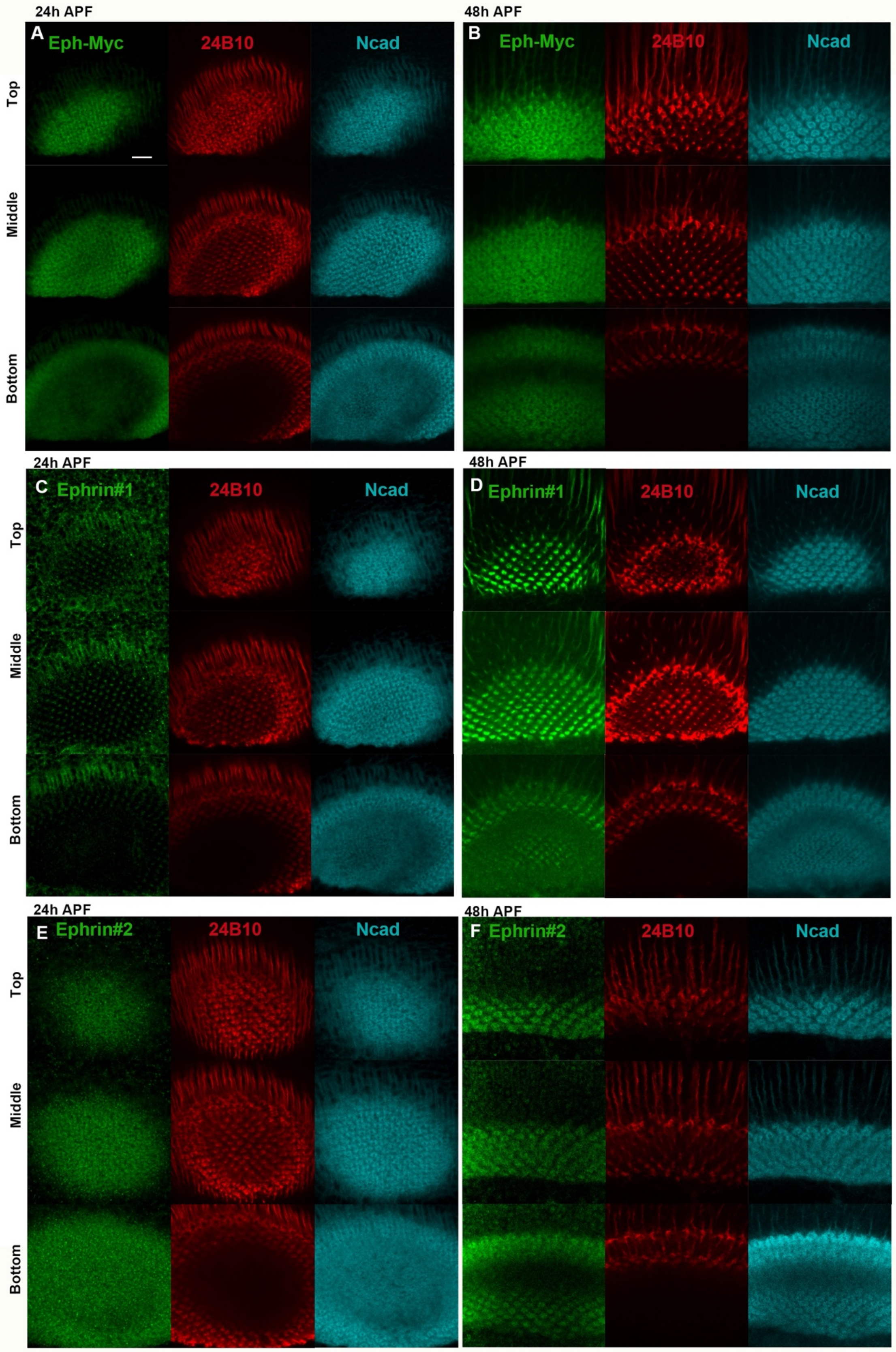

**Fig. S1. Eph and Ephrin distributions during pupal development.** (**A, B**) Eph-Myc shows a donut-like pattern (Myc, green) overlapping with Ncad (blue) at 24 h (**A**) and 48 h APF (**B**). (**C, D)** Ephrin #1 shows a dot-like pattern inside the donut-like domain of Ncad (blue) at 24 h (**C**) and 48 h APF (**D**). (**E, F**) Ephrin #2 shows a donut-like pattern overlapping with Ncad (blue) at 24 h and 48 h APF. 24B10 visualizes R7 and R8 terminals (red). Scale bars, 5 μm.

**Figure S2**

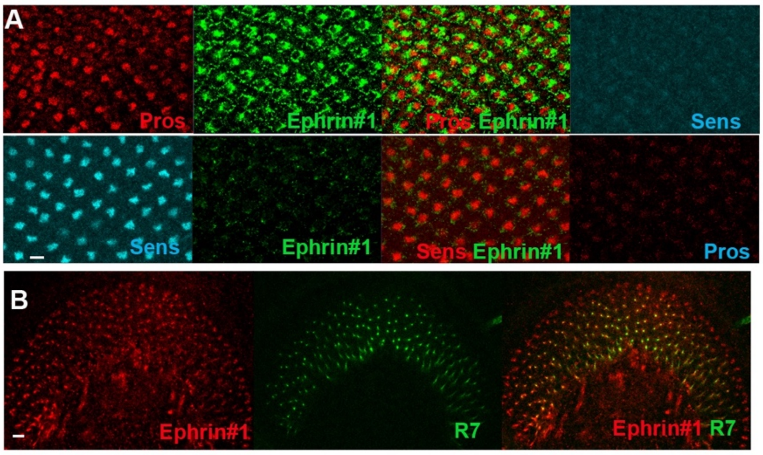


**Fig. S2. R7 is the source of Ephrin #1 signals.** (**A**) In the late third larval instar eye disc, strong Ephrin #1 signals are found in R7 cell bodies visualized by Pros (red), but not in R8 visualized by Sens (blue). (**B**) In the late third larval instar lamina, Ephrin #1 signals (red) overlap with R7 axons (green). Scale bars, 5 μm.

Figure S3

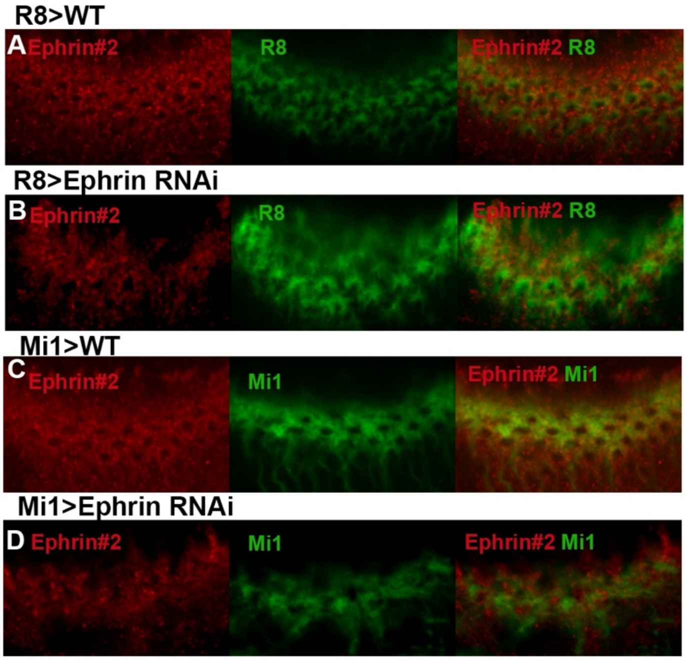


**Fig. S3. Ephrin#2 signals overlap with R8 and Mi1 terminals.** (**A-D**) *Ephrin* knock-down in R8 (**B**) and Mi1 (**D**) reduced Ephrin #2 signals (red) in R8 and Mi1 terminals (R8-GFP and Mi1-GFP, green), respectively. Scale bars, 5 µm.

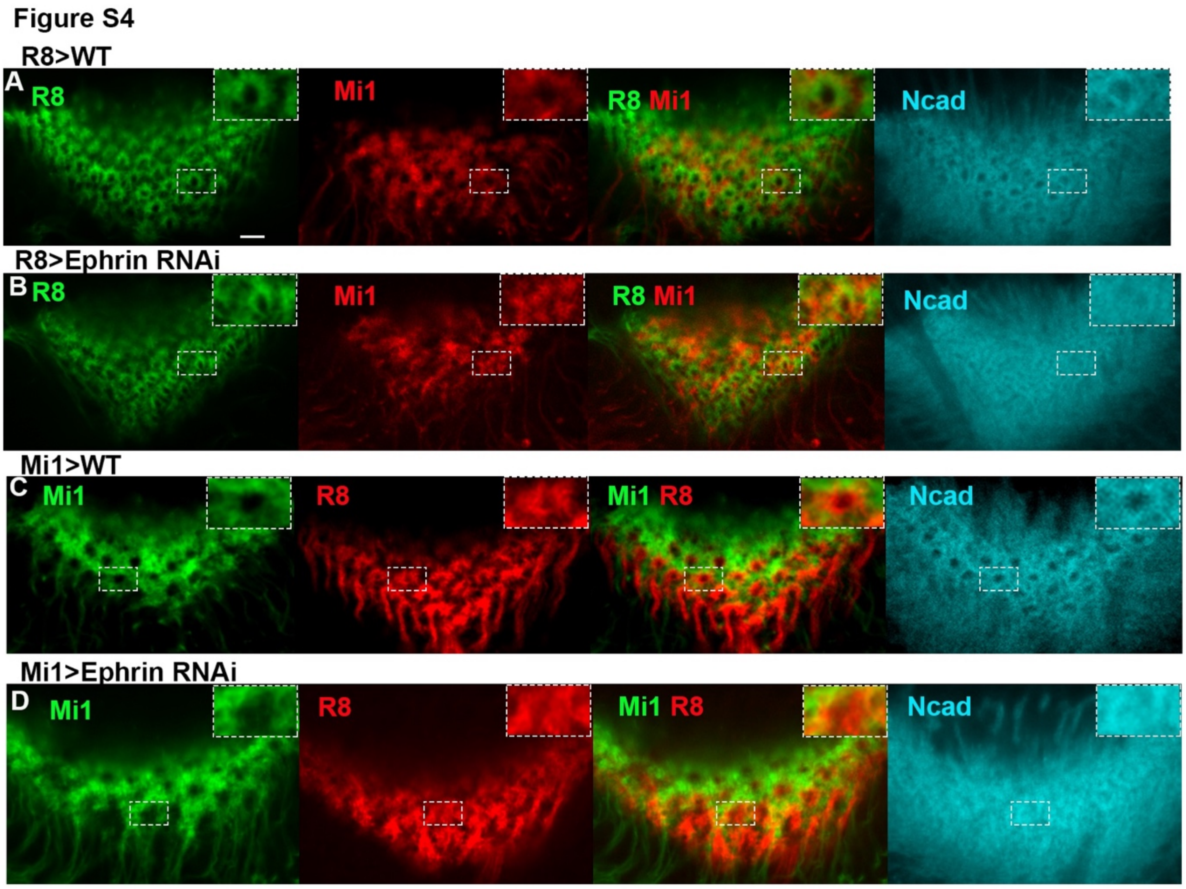


**Fig. S4. Ephrin expressed in R8 and Mi1 is non-autonomously required for column organization.** (**A, B**) R8 and Mi1 terminals (R8-GFP, green and Mi1-RFP, red) were disorganized upon *Ephrin* knock-down in R8. (**C, D**) R8 and Mi1 terminals (R8-GFP, red and Mi1-RFP, green) were disorganized upon *Ephrin* knock-down in Mi1. Scale bars, 5 µm.

Figure S5

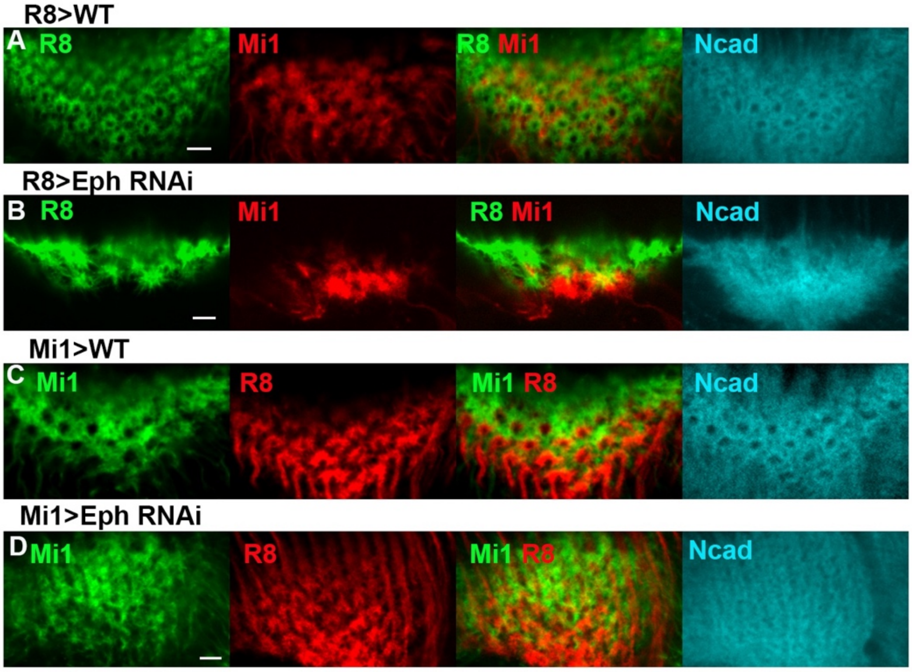


**Fig. S5. Eph expressed in R8 and Mi1 is required for column organization.** (**A, B**) R8 and Mi1 terminals (R8-GFP, green and Mi1-RFP, red) were disorganized upon *Eph* knock-down in R8. (**C, D**) R8 and Mi1 terminals (R8-GFP, red and Mi1-RFP, green) were disorganized upon *Eph* knock-down in Mi1. Scale bars, 5 μm.

**Figure S6**

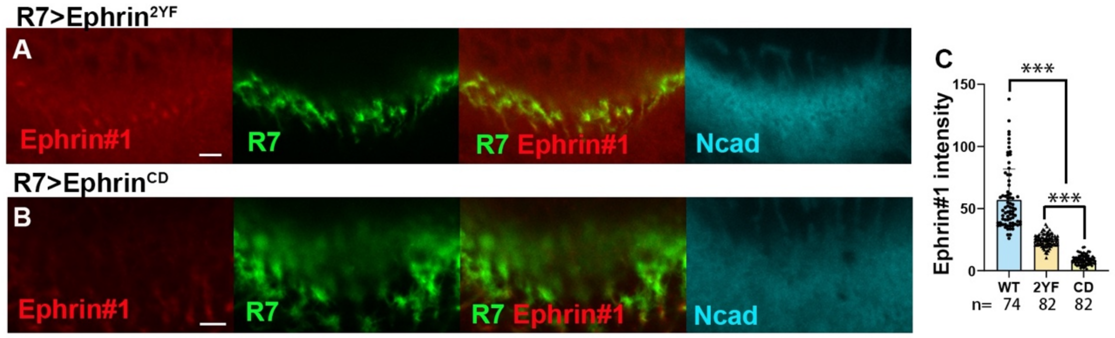


**Fig S6. Expression of mutant forms of *Ephrin* suppresses Ephrin phosphorylation.** (**A, B**) Ectopic expression of $Ephrin^{2YF}$ (**A**) and $Ephrin^{CD}$ (**B**) in R7 (R7-GFP, green) reduces Ephrin #1 signals (red). See Figure 2I as a control. (**C**) Quantification of Ephrin #1 intensity. *** indicates p< 0.001, n.s. not significant. Scale bars, 5 µm.

**Figure S7**

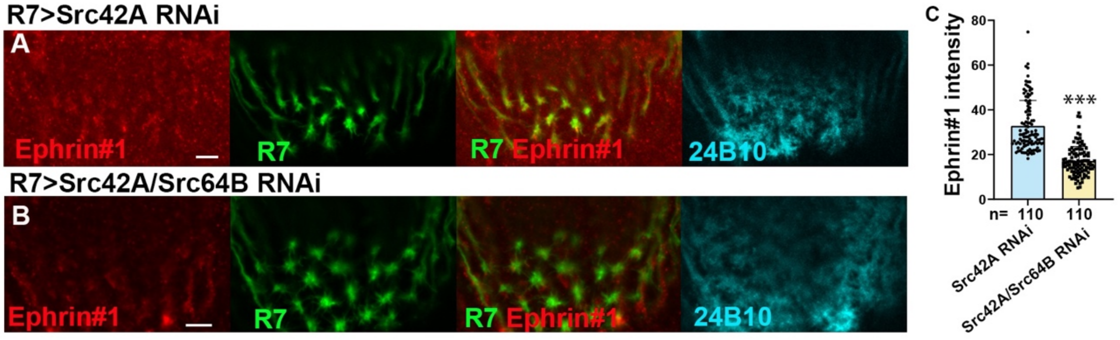


**Fig. S7. Src42A/Src64B double knockdown decreases Ephrin phosphorylation.** (**A**) *Src42A* knock-down in R7 (R7-GFP, green) reduced Ephrin #1 signals. See Figure. 2I as a control. (**B**) *Src42A* and *Src64B* double knock-down in R7 (R7-GFP, green) further reduced Ephrin #1 signals. (**C**) Quantification of Ephrin #1 intensity. *** indicates p< 0.001, n.s. not significant. Scale bars, 5 µm.
